# High-throughput screening of human genetic variants by pooled prime editing

**DOI:** 10.1101/2024.04.01.587366

**Authors:** Michael Herger, Christina M. Kajba, Megan Buckley, Ana Cunha, Molly Strom, Gregory M. Findlay

**Affiliations:** The Genome Function Laboratory, The Francis Crick Institute, London, United Kingdom; Viral Vector Core, Human Biology Facility, The Francis Crick Institute, London, United Kingdom

**Keywords:** prime editing (PE), multiplexed assay of variant effect (MAVE), *MLH1*, *SMARCB1*, saturation mutagenesis, functional genomics, genome editing technology, cancer predisposition

## Abstract

Understanding the effects of rare genetic variants remains challenging, both in coding and non-coding regions. While multiplexed assays of variant effect (MAVEs) have enabled scalable functional assessment of variants, established MAVEs are limited by either exogenous expression of variants or constraints of genome editing. Here, we introduce a pooled prime editing (PE) platform in haploid human cells to scalably assay variants in their endogenous context. We first optimized delivery of variants to HAP1 cells, defining optimal pegRNA designs and establishing a co-selection strategy for improved efficiency. We characterize our platform in the context of negative selection by testing over 7,500 pegRNAs targeting *SMARCB1* for editing activity and observing depletion of highly active pegRNAs installing loss-of-function variants. We next assess variants in *MLH1* via 6-thioguanine selection, assaying 65.3% of all possible SNVs in a 200-bp region spanning exon 10 and distinguishing LoF variants with high accuracy. Lastly, we assay 362 non-coding *MLH1* variants across a 60 kb region in a single experiment, identifying pathogenic variants acting via multiple mechanisms with high specificity. Our analyses detail how filtering for highly active pegRNAs can facilitate both positive and negative selection screens. Accordingly, our platform promises to enable highly scalable functional assessment of human variants.

## INTRODUCTION

Experiments to determine how genetic variants alter function can inform mechanisms, provide evidence for causal associations underlying human disease, and improve computational tools for variant effect prediction^1–3^. Attempts to leverage human genetic data for both biological discovery and precision medicine, however, have been hindered by a shortage of functional evidence. Despite improving performance of computational predictors^4–6^, rare variants are still highly challenging to interpret, both in coding and non-coding regions. Most variants observed in humans have never been assayed for functional effects, and hundreds to thousands of variants of uncertain significance (VUS) have been reported in each of many clinically actionable genes^7,8^.

Multiplexed assays of variant effect (MAVEs) have emerged as a means of providing functional evidence of variant effects at scale^9,10^. Key to their success is the ability to test many variants all in a single experiment. While numerous MAVEs have demonstrated high predictive power for identifying disease-associated variants^11–13^, technologies using genome editing to install variants have proven particularly accurate at detecting loss-of-function (LoF) variants in disease genes^14–16^, owing to advantages conferred by assaying variants in their native genomic context. For instance, endogenous levels of expression are maintained and variants with disruptive effects on both splicing and protein function can be readily identified.

However, current methods for assaying pools of variants via genome editing all have limitations. Saturation Genome Editing (SGE) leverages homology-directed DNA repair (HDR) to install hundreds of variants per experiment^17^. Yet, suboptimal scalability arises from low HDR rates in many cell types, as well as the requirement that variants be confined to a single region (100-200 bp) per experiment. Consequently, many separate SGE libraries are required to cover complete coding sequences^16^. Base editing screens are highly scalable and offer the advantage of being able to target sites genome-wide^18,19^, yet are limited by the fact that most substitutions cannot be made by a single base editor and by the potential of unintended editing at target sites to confound results.

We reasoned that prime editing (PE) systems may constitute a way forward by enabling virtually any short variant anywhere in the genome to be installed^20^. Prime editors are Cas9 nickases coupled to reverse transcriptase domains that create programmed edits via reverse transcription (RT). A single prime editing guide RNA (pegRNA) determines both the site of nicking and the variant to be edited into the genome. Original PE systems displayed lower efficiencies than evolved base editors used in recent screening applications^21^, but recent innovations have boosted the performance of PE in human cells. These include engineered prime editors for greater activity^22^, improved design of pegRNAs^23^, and manipulation of host repair pathways to increase desired PE outcomes^22^.

Whether these improved PE systems enable scalable and accurate functional characterization of variants genome-wide has yet to be established. Saturation Prime Editing (SPE) was the first implementation of PE for assaying variant libraries^24^. This work established that saturation mutagenesis of defined loci could be achieved with PE, and that variants could be assayed for functional effects with high accuracy. However, this required testing individual pegRNAs for activity prior to library design and direct sequencing of edited loci for effect quantification. More recently, PE screening using lentiviral delivery of pegRNAs was described, but effects of established pathogenic and benign variants were not readily distinguishable^25^. Therefore, while the potential of PE to introduce virtually any short variant at any locus in a cost-effective manner may prove ideal for testing large variant libraries, at present it remains unclear to what extent large PE screens are feasible. Challenges to overcome include, for instance, observing effects of variants that are recessive at the cellular level, and achieving sufficiently high editing rates such that variant effects can be accurately inferred from quantifying selection on pegRNAs.

Here, we set out to optimize a platform for engineering large numbers of variants with PE and to rigorously assess the platform’s potential to assay variants in both coding and non-coding regions, using both negative and positive selection. Our findings show that with stringent filtering for highly active pegRNAs, variants leading to functional effects can be accurately identified, suggesting that with further improvement and scaled implementation this platform will enable functional characterization of human variants genome-wide.

## RESULTS

### A pooled prime editing screening platform for efficient installation of diverse edits

We sought to develop a high-throughput screening platform that enables the functional interrogation of diverse genetic variants installed via prime editing (**Figure 1A**). To this end, we created a monoclonal HAP1 cell line with stable expression of the optimized prime editor PEmax^22^ via lentiviral integration (**Figure 1B**). HAP1 is a near-haploid human cell line with high value in genetic screening because recessive variant effects can be efficiently assayed^26^. Since HAP1 cells are mismatch repair (MMR) proficient, a second HAP1 line was generated with concomitant expression of a dominant negative MLH1 protein (MLH1dn) shown to enhance prime editing efficiency^22^.

**Figure 1.**
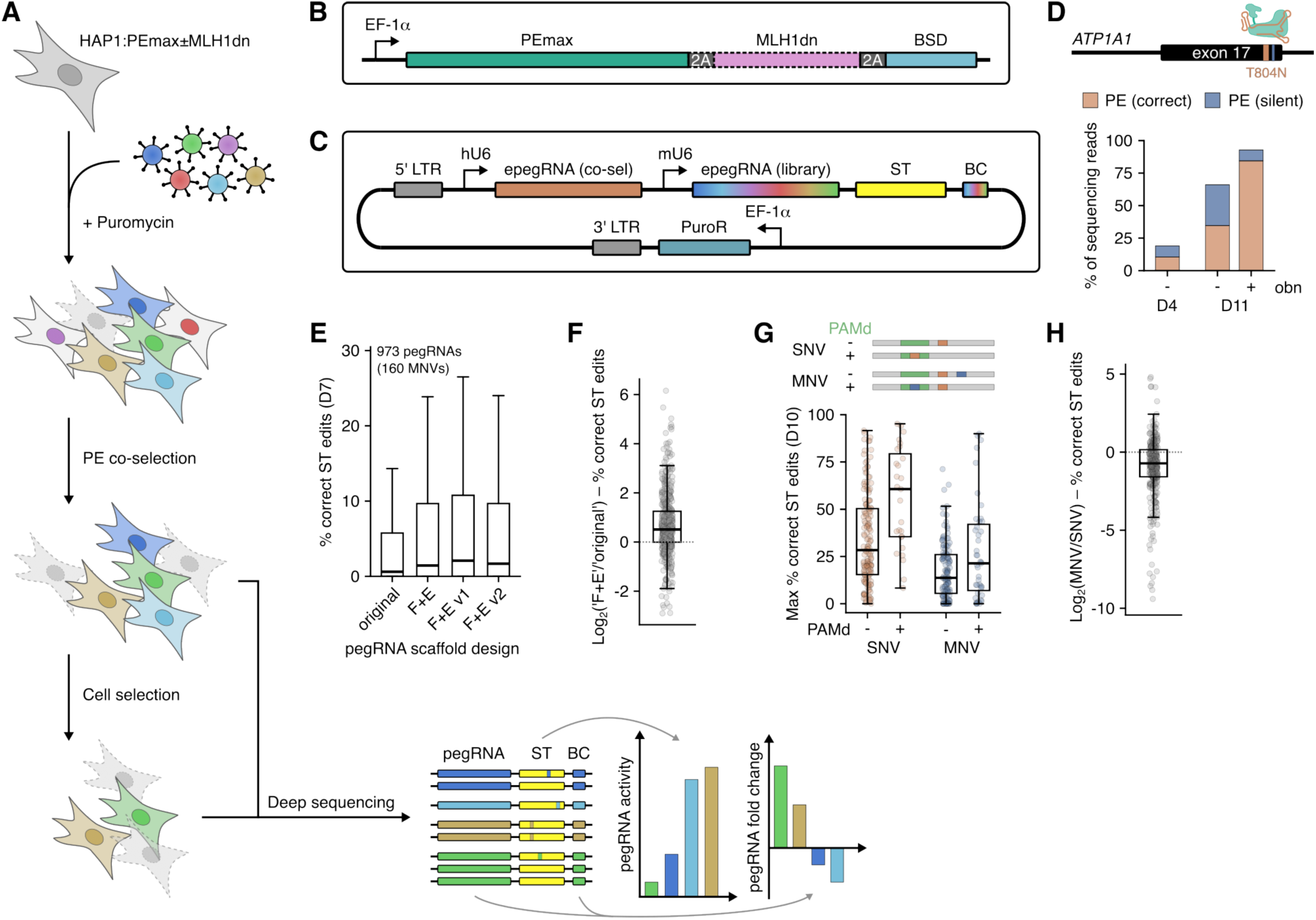
Development of a pooled PE screening platform in HAP1. (A) Schematic of the screening workflow, comprising lentiviral delivery of pegRNAs, puromycin selection, optional enrichment for edited cells via co-selection, functional selection (e.g. essentiality, drug treatment), deep sequencing of lentiviral cassettes, and quantification of pegRNA activities from surrogate targets (STs) and pegRNA frequency changes reflecting selective effects. Function scores for variants are determined from pegRNA frequency changes, using ST editing percentages to filter out inefficient pegRNAs. (B) Schematic of the construct stably integrated into the HAP1 genome for 2A-coupled expression of PEmax, MLH1dn, and the blasticidin S resistance cassette, expressed from a minimal EF-1α promoter. (C) Schematic of the vector for lentiviral delivery of pegRNA libraries. Each vector encodes a pegRNA to install a resistance mutation for co-selection and a pegRNA from the library to engineer a variant to be assayed. Each vector also includes a surrogate target (ST) matching the genomic sequence where the variant is to be introduced and a pegRNA-specific barcode (BC). (D) Enrichment of cells with the PE-mediated T804N edit in exon 17 of *ATP1A1* in response to ouabain (obn) treatment, as determined by amplicon sequencing. Percentages of reads with correct PE (T804N) or partial PE (silent mutation only) are shown. (E) Boxplot of ST editing percentages for 973 pegRNAs using different scaffolds (bold line, median; boxes, interquartile range (IQR); whiskers extend to points within 1.5x IQR; outliers not shown). (F) Boxplot of log2-fold-changes in ST editing percentages per variant using pegRNAs with the F+E scaffold compared to matched pegRNAs with the original scaffold (data from E). (G) Schematic of SNV and MNV types showing target edit (orange) and additional silent mutation (blue). Boxplot of correct ST editing percentages for the top pegRNA per variant grouped by edit type and PAM disruption (PAMd). (H) Boxplot of log2-fold-changes in ST editing percentages comparing MNV- and SNV-pegRNA pairs (data from G).

We reasoned that akin to other pooled CRISPR screening modalities^18,19,27^, genomic integration and deep sequencing of pegRNAs over time could serve as a functional readout, provided each pegRNA efficiently introduces the variant it encodes. Such a system would afford highly scalable interrogation of variant effects without being confined to a single locus or gene per experiment.

Towards optimizing PE efficiencies, we leveraged engineered pegRNAs (epegRNAs)^23^. Specifically, we incorporated the structured RNA motif tevopreQ_1_ as a stabilizing 3’ extension cap. Hereafter, epegRNAs modified in this manner will be referred to simply as pegRNAs.

pegRNA design tools such as PRIDICT and DeepPrime^28,29^ now offer predictions of editing efficiency for user-defined variants. However, our work predates these models, so we designed pegRNAs using PEGG^30^. To maximize our chances of including an efficient pegRNA for each variant, in our libraries we allowed up to nine different pegRNA designs per variant, with diverse spacer and 3’ extension sequences.

We reasoned that an integrated readout of pegRNA activity may be important for accurate variant scoring due to the large variability of PE efficiencies within pegRNA pools. Therefore, downstream of each pegRNA, we included 55 bp of genomic target sequence to serve as a surrogate target (ST), capable of informing pegRNA editing efficiency after genomic integration (**Figure 1C**). STs have been previously employed for pooled base editing^31^ and (pe)gRNA activity screens^32^.

We experimentally tested the following strategies to further optimize variant installation via PE: 1) enrichment via co-selection, 2) use of optimized pegRNA scaffolds, and 3) incorporation of PAM-disruptive synonymous edits in pegRNA design. Co-selection systems enable the enrichment of intended edits by co-editing a second locus leading to a selectable phenotype. This strategy has been shown to increase rates of intended edits in NHEJ- and HDR-mediated editing^33^, as well as in base editing and prime editing^34,35^. In our system, we adopted a PE co-selection strategy in which a SNV within *ATP1A1*, a HAP1-essential gene, leads to resistance against the Na^+^/K^+^- ATPase inhibitor ouabain^35^.

First, we identified a pegRNA which efficiently installs T804N with a silent, PAM-disruptive mutation in HEK293T cells (**Figure S1**). We next determined editing and ouabain selection performance with this pegRNA when expressed stably in our HAP1 PE lines, observing 10.4% of cells with the intended T804N mutation at the earliest timepoint post-transduction (**Figure 1D**). Continued culture of cells for 7 more days in the absence of ouabain resulted in a 3.3-fold increase in T804N editing to 34.6%, indicating that stable expression of all PE components leads to continuous accumulation of edits. After 7 days of ouabain selection, 84.4% of alleles contained both T804N and the silent PAM mutation. This experiment validates efficient *ATP1A1* editing and ouabain selection in our HAP1-based PE system, enabling assessment of ouabain-based co-selection during screening.

Structurally stabilized guide RNA (gRNA) scaffolds have been shown to enhance various Cas-based activities^36–38^. We compared the performance of four different scaffold designs with a set of 973 pegRNAs installing 160 multi-nucleotide variants (MNVs) in *SMARCB1* (up to 9 pegRNAs per variant). This experiment was conducted via lentiviral delivery of pooled pegRNA libraries to HAP1:PEmax+MLH1dn. Editing rates for each pegRNA-scaffold combination were assessed 7 days post-transduction via deep sequencing of pegRNA-ST cassettes to quantify the percentage of ST reads containing correct edits (**Supplementary Table 1**). Incorrect pegRNA-ST cassettes (e.g., due to recombination or errors in synthesis or sequencing) were discarded from analysis (see **Methods**, **Figure S2**).

pegRNAs with the original gRNA scaffold^39^ achieved a median editing rate of 0.6% (**Figure 1E**). All stabilized scaffolds performed better than the original. The F+E scaffold, comprising an A-U flip and stem extension^36^, achieved a 2.4-fold improvement in median PE efficiency over the original scaffold (**Figure 1F**). 87% of all pegRNAs exhibited higher or equal editing activity using the F+E scaffold. Swapping the flipped A-U bases in the F+E scaffold with a G-C pair (v1)^37^ lead to a comparable increase in editing (median 2.1% correct ST editing; **Figure 1E**). Further stabilization by introducing a superstable loop within the first hairpin of the scaffold (v2), as reported in the t-lock design^38^, did not increase pegRNA activities. In summary, stabilized pegRNA scaffold designs proved optimal for obtaining maximal PE efficiency across a large number of pegRNAs.

Next, we investigated whether SNVs are most efficiently introduced as individual mutations or with additional edits. Previous work has suggested that programming silent mutations in addition to each target SNV may be advantageous because MNVs are less efficiently recognized by the MMR machinery^22^. To test this in our platform, we designed two pegRNA libraries, one encoding 191 distinct SNVs on their own and the other programming the same set of SNVs with 1 or 2 additional silent mutations, creating 166 corresponding MNVs in total. The libraries were pooled to a total of 2,188 pegRNAs and assayed in HAP1:PEmax+MLH1dn to measure ST editing rates for each pegRNA on day 10 (**Supplementary Table 2**).

Only the best-performing pegRNAs per variant were analyzed to account for differences in how restrictive pegRNA design may be for either SNVs or MNVs. It is established that mutations within the protospacer adjacent motif (PAM) are more efficiently installed by PE and, indeed, we found that PAM-disruptive edits achieved 2.1- or 1.6-fold higher editing as SNVs or MNVs, respectively (**Figure 1G**). Overall, however, we observed higher editing rates for SNVs than MNVs independent of PAM disruption (median 61% increase in editing comparing SNVs to MNVs; **Figure 1H**). As most SNVs were installed more efficiently as single substitutions, we proceeded with screening libraries designed in this manner.

### High-throughput essentiality screening of *SMARCB1* variants

We next analyzed whether our pooled PE platform would enable the functional interrogation of variants via negative selection (**Figure 2A**). Specifically, we asked whether sequencing of pegRNA-ST cassettes alone could accurately define variants with deleterious effects (i.e. without needing to perform endogenous target site sequencing).

**Figure 2.**
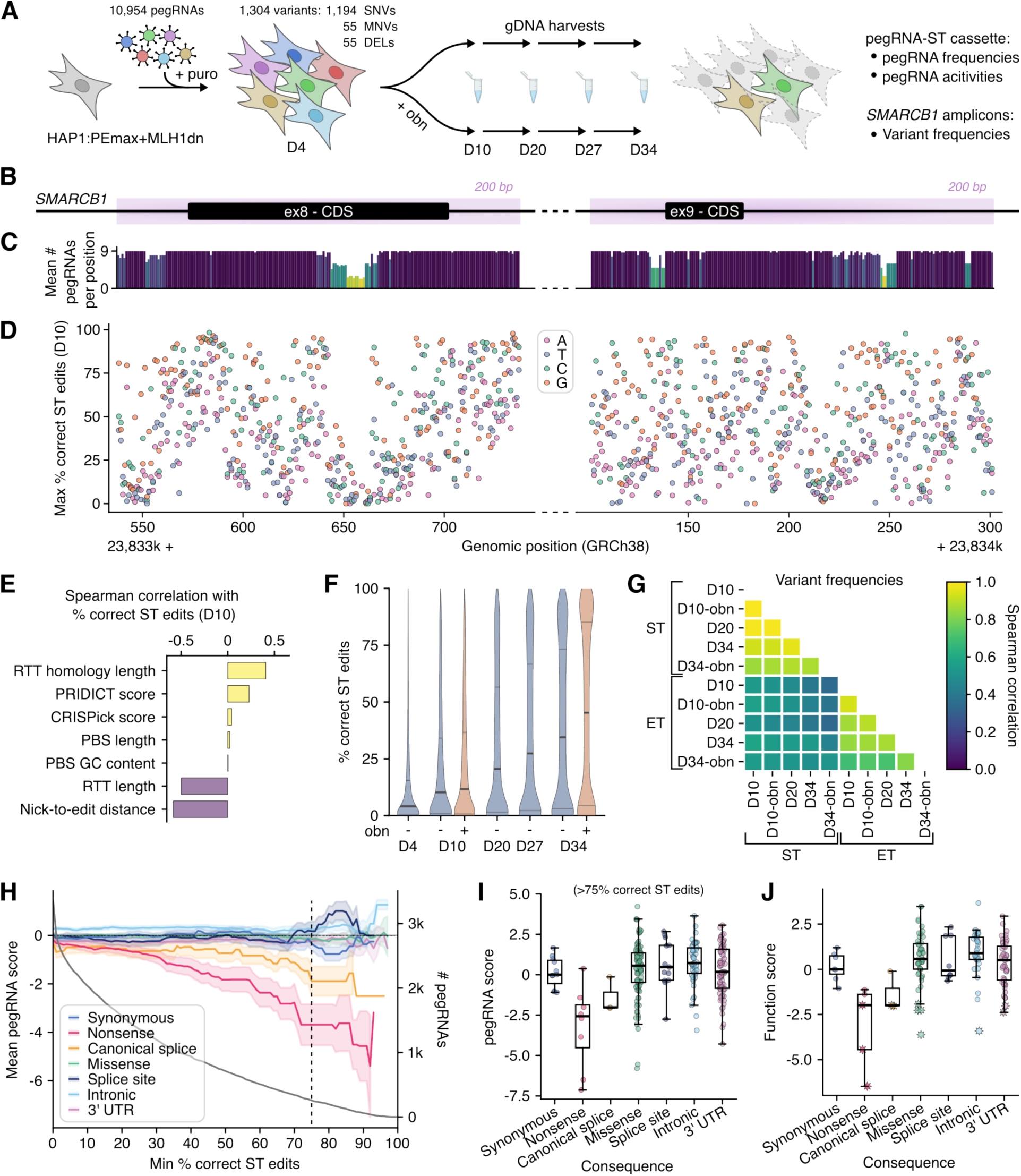
Near saturation mutagenesis of *SMARCB1* regions by pooled PE. (A) Schematic of the experimental workflow for *SMARCB1* essentiality screening using pooled PE, comprising lentiviral delivery of pegRNAs to HAP1:PEmax+MLH1dn cells (D0), and culture until D34 with sequential harvesting for deep sequencing of pegRNA-ST cassettes and *SMARCB1* target regions. (B) Illustration of target regions (light purple) for saturation mutagenesis, centered on *SMARCB1* exons 8 and 9. (C) For each genomic position, the average number of pegRNAs per SNV included in the library is plotted. pegRNA coverage per variant was capped at 9. (D) The maximum ST editing rate for each SNV 10 days post-transduction is plotted by position. (E) Spearman correlation of correct ST editing percentages with different pegRNA features for a pool of 8,612 pegRNAs programming SNVs, MNVs and deletions (DELs) across *SMARCB1*. ST editing percentages were averaged across D10 duplicates without ouabain before correlation analysis. (F) Violin plots of ST editing percentages observed across the pegRNA pool at multiple timepoints, with and without ouabain (obn) co-selection. Bold line indicates the median and dashed lines the interquartile ranges. (G) Heat map of Spearman correlation coefficients determined between surrogate target (ST) and endogenous target (ET) variant frequencies across samples. ST variant frequencies were computed as the fraction of all STs containing a given variant. ET variant frequencies were adjusted with a variant-level background-correction based on negative control sample sequencing. (H) Mean log2-fold-change in pegRNA frequency (D34 over D10) in ouabain-treated samples, by variant consequence. Mean values are plotted as a function of ST editing thresholds used for pegRNA filtering. Bands correspond to 95% confidence intervals, and the gray curve shows the number of pegRNAs passing frequency and ST editing thresholds. The dashed vertical line corresponds to 75% correct ST edits, chosen as the threshold for analysis in (**I** and **J**). pegRNAs with frequencies lower than 6×10^-5^ on D10 were excluded from analysis. (**I**) pegRNA log2-fold-change between D34-ouabain and D10-ouabain samples, grouped by consequence. (**J**) pegRNA scores from (**I**), averaged per variant (function scores). Significantly scored variants (FDR<0.05) are indicated with stars. (Boxplots: bold line, median; boxes, IQR; whiskers extend to points within 1.5x IQR).

*SMARCB1* encodes a core subunit of the BAF (BRG1/BRM-associated factor) chromatin remodeling complex and inactivating mutations are known to drive diverse cancers^40^. *SMARCB1* is also among ∼2,200 essential genes in HAP1^41^, meaning LoF variants should be depleted in culture. We independently confirmed *SMARCB1* essentiality in HAP1 by observing depletion of frameshifting indels after Cas9-mediated editing (**Figure S3**).

Exons 8 and 9 of *SMARCB1* fully encode the C-terminal coiled-coil domain (CTD), which harbors recurrently observed missense mutations in cancers and the rare developmental disorder Coffin- Siris syndrome^40,42^. We designed a pegRNA pool to install all possible SNVs in two 200-bp regions across exons 8 and 9 (**Figure 2B**). Additionally, we included MNVs encoding nonsense mutations as well as 3-bp deletions of every codon within the target regions.

We programmed variant-level redundancy in the pegRNA library, allowing a maximum of 9 pegRNA designs per variant, which was possible for most SNVs (**Figure 2C**). Although pegRNA coverage for some mutations was lower due to the scarcity of NGG-PAM sequences, we successfully designed pegRNAs for nearly all intended SNVs (99.5%), MNVs (100%) and deletions (100%).

In total, 10,954 pegRNAs programming 1,194 SNVs, 55 MNVs, and 55 deletions were cloned into a lentiviral expression vector with the F+E scaffold. In addition to STs, the cassette included the co-selection pegRNA (*ATP1A1*-T804N) for optional enrichment of edited cells. The resulting library was introduced to HAP1:PEmax+MLH1dn via lentiviral transduction (multiplicity of infection (MOI) less than 0.5) and deep sequencing of pegRNA-ST cassettes was performed on days 4, 10, 20, 27, and 34 to measure pegRNA frequencies and editing dynamics. To test the co-selection strategy, one pool of cells was treated with ouabain on day 5, and two replicate pegRNA libraries without co-selection were also maintained (see **Methods**). We observed strong correlations for pegRNA frequencies and ST editing percentages between replicate libraries and across timepoints (**Figures S4A and S4B**).

We assessed PE efficiency of pegRNAs designed to introduce SNVs across both target regions (exons 8 and 9). First, to assess variant-level editing activity from sequencing of pegRNA-ST cassettes, ST editing percentages were averaged across all pegRNAs encoding the same edit and mapped to the genomic coordinate of each variant. This revealed highly variable PE efficiencies within the pegRNA pool (**Figure S4C**). 40% of pegRNAs had ST editing rates above 10%. We also checked the top-performing pegRNA per SNV and found that 89.9% of all SNVs were installed with at least 10% ST editing by at least one pegRNA (**Figure 2D**). Editing rates across target regions were highly variable, except for a few regions of lower PE efficiencies that coincided with low pegRNA coverage.

We next asked which pegRNA features correlated with higher PE efficiencies. Of pegRNA features analyzed, RTT homology length was most strongly correlated with ST editing on day 10 (R = 0.43, r = 0.37), followed by PBS length and PBS GC-content (**Figure 2E**). On the contrary, total RTT length and nick-to-edit distance were negatively correlated with ST editing. We also observed a modest correlation with CRISPick scores (i.e., predictions of gRNA on-target activity)^37,43^. Finally, we retrospectively computed PRIDICT scores^28^ where possible for our pegRNAs, and found a reasonable correlation with experimentally determined activities, demonstrating the value of this tool for future pegRNA design (**Figures S4D and S4E**).

One potential advantage of stably expressing all PE components is that intended edits may accumulate in culture over time. Indeed, for all libraries tested we observed a continued increase in ST editing with prolonged cell culture (**Figure 2F**). Median ST editing rates across the pegRNA pool increased from a modest 4.1% on day 4 to 34% on day 34 for samples without co-selection. Overall ST editing was highest with the addition of ouabain-mediated co-selection on day 5, reaching a median efficiency of 45% by day 34. This was a modest improvement compared to samples without co-selection. Notably, ST editing percentages across all samples were bimodal by day 34. For instance, with co-selection, 33.0% of pegRNAs showed greater than 75% editing, while 30.6% of pegRNAs showed less than 10% editing.

For optimal scalability, it would be ideal if the presence of a variant in the genome could be inferred from sequencing of the pegRNA-ST cassette alone, but for this to work effectively, ST editing rates must accurately reflect endogenous target (ET) editing rates. To assess this, we sequenced ETs in *SMARCB1* amplified from the same cells used to measure pegRNA frequencies and ST editing. In the ouabain co-selected day 10 sample, we identified 79% of all intended variants at frequencies greater than background (defined via sequencing of unedited controls). Because multiple pegRNAs were used to engineer each variant, we used the variant-level ST editing rate calculated for each set of pegRNAs encoding a single variant for comparison to endogenous variant frequencies (**Figures 2G and S4F**), observing a correlation of r = 0.53 for all variants, and r = 0.69 for MNVs and DELs. This demonstrates ST editing can be used as a reasonable proxy for identifying pegRNAs that install variants with high activity.

Observing that nearly all variant frequencies in ETs slowly increased during the 34-day experiment, it was unclear whether selection against pegRNAs encoding LoF variants could be determined from sequencing pegRNAs alone (**Figures S4G and S4H**). To investigate this, we calculated a pegRNA score for each pegRNA in the library, defined as its day 34 frequency normalized to its day 10 frequency (**Supplementary Table 3**). We also calculated variant-level function scores by averaging scores from pegRNAs installing the same variant (**Supplementary Table 4**).

Given selection is predicted to be strongest against the most active pegRNAs encoding LoF variants, we explored how different ST editing activity filters impact our ability to separate signal from noise. If no threshold is applied, or only a lax threshold set to 10% ST editing on day 10, there is no clear separation between pegRNAs installing nonsense variants and synonymous variants (**Figure 2H**). However, using an ST editing threshold of 30%, pegRNAs encoding nonsense and canonical splice variants score distinctly lower than pegRNAs encoding synonymous variants. This trend becomes more pronounced if only highly active pegRNAs are retained, with a mean nonsense pegRNA score of −3.70 when using a stringent ST editing threshold of 75%. While effects of negative selection are readily apparent for highly active pegRNAs, imposing a stringent threshold excludes most pegRNAs tested, resulting in a considerable reduction in the number of variants successfully assayed (**Figure 2H**). Nevertheless, applying an ST editing threshold of 75% enables many variants predicted to be LoF (i.e., nonsense, canonical splice) to be distinguished by low pegRNA scores and low function scores (**Figures 2I-J and S5**).

Among 164 variants scored with this filter, we identified 12 significantly depleted variants, including 3 nonsense and 2 canonical splice variants, using a false discovery rate (FDR) of 0.05 to call LoF variants (see **Methods**). LoF missense variants clustered within a highly conserved ⍺- helix of the SMARCB1 CTD. We also scored the intronic SNV c.1119-12C>G as LoF, which is annotated as a VUS in ClinVar^7^ but has been hypothesized to disrupt splicing^44^. Indeed, our functional data and a SpliceAI score^4^ of 0.96 corroborate this effect. Overall, these examples illustrate how select LoF variants efficiently installed by PE can be functionally identified via negative selection and suggest that further improvements to editing efficiency will enable many more variants to be assessed in this manner.

### Accurate determination of variant effects on *MLH1* function

We next used our platform to assay variants across *MLH1*, a tumor suppressor gene encoding a subunit of the MMR pathway. Germline pathogenic variants in *MLH1* predispose patients to several types of cancer, including colorectal and endometrial carcinoma, and nearly 2,000 VUS in *MLH1* have been reported in ClinVar^7^. Unlike *SMARCB1*, *MLH1* is not essential in HAP1. However, it has been shown that loss of MMR pathway function in HAP1 leads to partial 6- thioguanine (6TG) resistance^45^. Therefore, 6TG selection may enable positive selection of pegRNAs encoding LoF variants. One advantage of this approach is that selection can be initiated after an extended period of editing, therefore increasing the fraction of edited cells at the onset of selection.

To assess 6TG selection, we performed dilution series in wildtype (WT) HAP1, HAP1:PEmax+MLH1dn, and HAP1:PE2+MLH1-knockout (KO) cells (**Figure S6A**). Compared to WT, HAP1:PE2+MLH1-KO cells were highly resistant to low doses of 6TG, whereas HAP1:PEmax+MLH1dn cells were only mildly resistant. Reasoning that use of HAP1:PEmax+MLH1dn cells for screening may lead to more efficient prime editing but less efficient selection, we performed screens in both HAP1:PEmax and HAP1:PEmax+MLH1dn using different 6TG concentrations (1.2 µg/ml and 1.6 µg/ml, respectively). Screens were also performed both with and without ouabain co-selection.

Our first screen aimed to identify LoF SNVs in exon 10 of *MLH1*. The exon 10 library consisted of 2,696 pegRNAs encoding 598 variants, including nearly all possible SNVs (96%) in a 200-bp region spanning exon 10 and flanking intronic regions, as well as 22 nonsense MNVs (**Figure 3A**). Libraries were designed, cloned, transduced, and sequenced as for *SMARCB1* experiments, with genomic DNA (gDNA) samples collected on days 4, 13, 20, and 34. Ouabain co-selection was performed from day 13 to day 20, and 6TG selection was initiated on day 20 prior to harvesting surviving cells on day 34. Sequencing of both pegRNA-ST cassettes and endogenous loci was performed.

**Figure 3.**
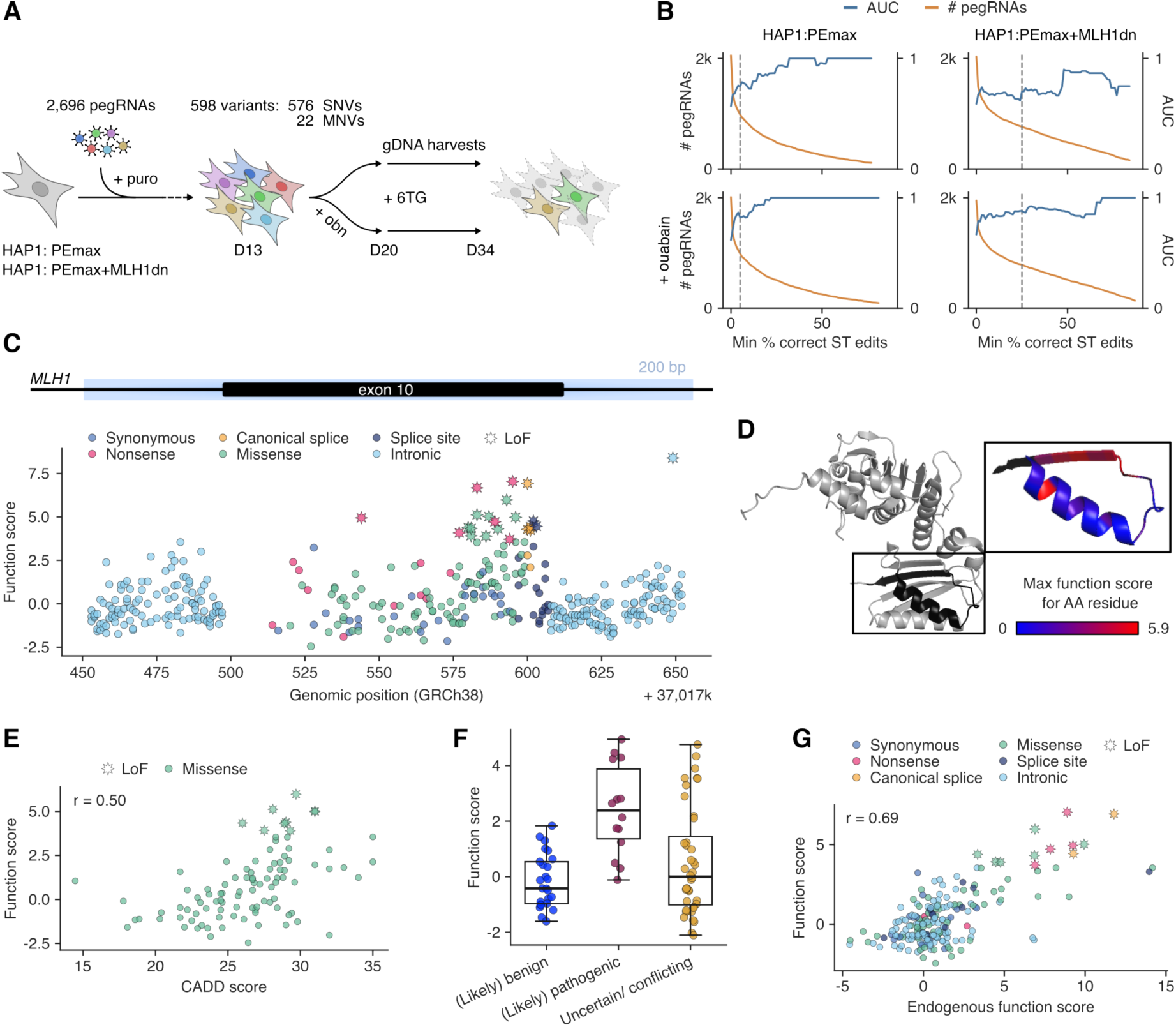
Pooled prime editing of *MLH1* exon 10 identifies disease-associated variants. (A) Experimental workflow of *MLH1* 6TG selection screens, comprising lentiviral delivery of pegRNAs to HAP1:PEmax and HAP1:PEmax+MLH1dn cells, ouabain co-selection (D13-D20), cell harvesting before (D20) and after (D34) 6TG challenge, and deep sequencing of pegRNA-ST cassettes and the *MLH1* target region. (B) For each condition, AUC measurements for pLoF variant identification are plotted as a function of ST editing threshold. For this analysis, synonymous variants were defined as pNeut and nonsense and canonical splice variants as pLoF. (C) Function scores for *n* = 401 variants are plotted by genomic position and colored by variant consequence. Significantly scored variants (q-value less than 0.01) are indicated with stars. (D) The highest function score of all missense variants assayed at each amino acid position is shown on the *MLH1* structure (PDB: 4P7A). (E) Correlation between function scores and CADD scores for *n* = 98 missense variants (Pearson’s r = 0.50). (F) Function scores of *n* = 79 variants present in ClinVar, grouped by pathogenicity annotation. (G) Correlation between function scores (pegRNA-derived) and endogenous function scores for *n* = 205 variants.

We first compared ST editing and ET editing across cell lines and co-selection strategies. We observed higher editing at both STs and ETs in the HAP1:PEmax+MLH1dn cell line compared to HAP1:PEmax (**Figure S6B**). In both lines, ouabain co-selection modestly improved editing at day 20. For instance, the median ST editing rate on day 20 was 11.9% in ouabain-treated HAP1:PEmax+MLH1dn cells, compared to 9.1% in cells without ouabain, and in the same line, the median variant frequency in ET sequencing increased from 2.4x10^-4^ to 3.0x10^-4^ with addition of ouabain. As before, editing rates of individual pegRNAs were highly variable. For instance, 40% of pegRNAs had ST editing percentages over 25% in ouabain-treated HAP1:PEmax+MLH1dn cells.

To investigate whether LoF variants could be identified from changes in pegRNA frequencies over time, we calculated pegRNA scores as the log2-ratio of day 34 pegRNA frequency over day 20 pegRNA frequency (**Supplementary Table 5**). As before, function scores for individual variants were obtained by averaging scores of all pegRNAs encoding the same variant with ST editing rates greater than a specified threshold (**Supplementary Table 6**). To assess performance for identifying LoF variants, we defined synonymous variants as putatively neutral (pNeut) and nonsense and canonical splice variants as putatively LoF (pLoF), then calculated area under the receiver operating characteristic curve (AUC) measurements using a continuous range of ST editing thresholds. This approach revealed thresholds that enable accurate detection of pLoF variants across all 4 screening conditions (**Figure 3B**). AUCs reached 1.00 in 3 of 4 screens using more stringent thresholds. For example, an AUC of 1.00 was reached using an ST editing threshold of 22% for pegRNAs assayed in HAP1:PEmax with co-selection (**Figure 3B**), while maintaining 20% of pegRNAs designed.

To include more pegRNAs in analysis, relatively lax ST editing thresholds were set at 5% for experiments in HAP1:PEmax, and at 25% for experiments in HAP1:PEmax+MLH1dn. While more stringent filters may perform optimally for a single condition, this approach allowed more pegRNA scores to be compared across screen conditions (**Figure S7A**).

We assessed how well pegRNA scores and function scores were correlated between conditions, both before and after filtering pegRNAs on ST editing rates (**Figure S8A-C**). Correlations of pegRNA scores improved with filtering, in turn resulting in well correlated function scores. To determine a final function score for each variant, we required the variant to have been assayed in at least two conditions. This approach led to function scores for 401 out of 598 variants for which pegRNAs were designed, determined from 967 retained pegRNAs.

To define LoF variants, we defined an empirical null distribution from synonymous variants and set the FDR to 0.01 (**Methods**). Mapping function scores to their genomic position reveals a cluster of LoF missense variants near the end of exon 10 (**Figure 3C**). These missense variants localize to a highly conserved β-sheet in the MLH1 structure (PDB: 4P7A, **Figure 3D**), revealing residues intolerant to mutation.

To orthogonally validate our exon 10 results, we compared function scores for missense variants to CADD scores, observing a Pearson’s correlation of r = 0.50 (**Figure 3E**). As many variants in exon 10 have been reported in ClinVar, we also compared function scores across pathogenicity categories. While 57% of “pathogenic” and “likely pathogenic” variants in the region had function scores greater than 2.0, no “benign” or “likely benign” variants scored over 2.0 (**Figure 3F**). This suggests our platform may enable scalable identification of new pathogenic variants. Indeed, 7.5% of VUS tested in this region were LoF in our assay.

Depending on intended use, more stringent filtering of pegRNAs based on ST editing may be implemented. For example, we repeated analyses only on variants with average ST editing rates greater than 36%, corresponding to the lowest threshold at which AUC = 1.00 for distinguishing pLoF from pNeut variants. This approach improved the correlation to CADD scores (r = 0.64) and resulted in non-overlapping score ranges for *n* = 5 pathogenic SNVs and *n* = 9 benign SNVs (**Figure S9**).

Finally, we performed sequencing of the endogenous *MLH1* exon 10 region to quantify variants’ enrichment in genomic DNA following selection (**Figures 3G and S10A-B**). The highest scoring variants, as determined by pegRNA-based function scores, were also highly enriched in endogenous DNA. The correlation between function score and endogenous function score (defined as the log2-ratio of post-6TG frequency over pre-6TG frequency for each variant in ET sequencing) increased from 0.68 to 0.77 when analyzing only variants with average ST editing percentages greater than 36%. In summary, these experiments demonstrate successful identification of LoF variants with relatively high SNV coverage in a single genomic region, while once more highlighting the importance of stringent pegRNA filtering.

### Screening all short non-coding *MLH1* variants in ClinVar in a single experiment

Non-coding variants are challenging to interpret clinically and have proven difficult to assay at scale via genome editing, especially when distributed over large sequence spaces. To evaluate whether our PE platform could be used for high-throughput assessment of non-coding variants, we designed a pegRNA library encoding all non-coding *MLH1* variants smaller than 10 bp and reported in ClinVar. This library consisted of 3,748 pegRNAs covering 874 variants distributed across 60 kb of *MLH1*, with many variants located near intron-exon boundaries (**Figure 4A**).

**Figure 4.**
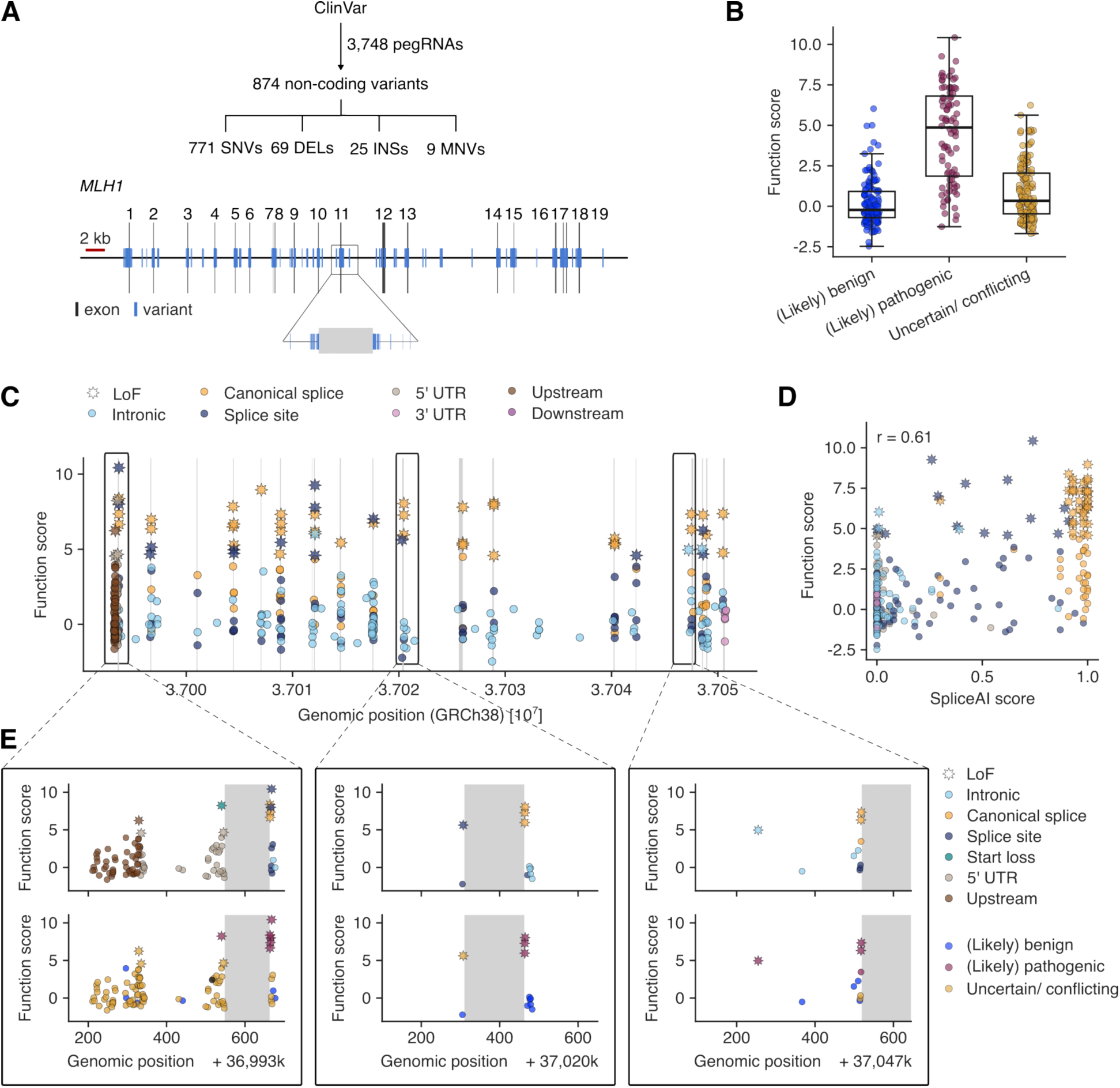
Screening all non-coding *MLH1* variants in ClinVar for LoF effects. (A) A library of 3,748 pegRNAs was designed to install 874 non-coding variants from ClinVar including 771 SNVs, 69 deletions (DELs), 25 insertions (INSs), and 9 MNVs. Variants, represented by blue lines, span the entire *MLH1* locus. (B) Function scores are plotted for *n* = 357 variants, grouped by pathogenicity annotation in ClinVar. (C) Function scores are plotted for *n* = 362 variants by genomic position and color-coded by variant consequence. Stars indicate variants with *q*-values less than 0.01. (D) The correlation between function scores of SNVs and spliceAI scores is plotted (*n* = 296). (E) Detailed results for select regions are shown: exon 1 and upstream (left), exon 11 (middle), and intron 15 (right). Exonic regions are in gray, and variants are colored by consequence (upper panels) or ClinVar annotation (lower panels).

Using the same strategy as for the *MLH1* exon 10 experiment, including repeating screens across four conditions and filtering pegRNAs by activity, we derived final function scores for 362 of the 874 variants tested (**Supplementary Tables 7,8**). Compared to the results for exon 10, we observed modestly improved correlations of pegRNA scores and function scores across conditions (**Figures S8D-F**). Once more, filtering of pegRNAs by ST editing was critical to see clear differences between pLoF variants and pNeut variants, defined for this experiment as canonical splice site variants and intronic variants more than 8 bp from any exon, respectively (**Figure S7B**). The screen that best discriminated pLoF variants was performed in HAP1:PEmax with ouabain treatment (**Figure S11**), again confirming an advantage of co-selection.

Importantly, the vast majority of ClinVar-pathogenic variants assayed had function scores greater than 1.5 (median = 4.87, S.D. = 2.90; **Figure 4B**). In contrast, nearly all benign variants scored neutrally (median = −0.22, S.D. = 1.45), while scores of VUS spanned a wide range (median = 0.34, SD = 1.72). Using a stringent threshold to determine LoF variants (*q* < 0.01; **Figure 4C**), we called 54% of ClinVar-pathogenic variants as LoF and only 2.4% of ClinVar-benign variants as LoF (AUC = 0.89 for distinguishing pathogenic and benign).

As intronic variants deemed LoF are likely to impact splicing, we compared function scores to SpliceAI scores for orthogonal validation (**Figure 4D**). While the highest scoring variants by SpliceAI tended to have high function scores, many variants with intermediate SpliceAI scores (0.25-0.75) also scored as LoF. Overall, only 2 of *n* = 96 intronic variants with SpliceAI scores less than 0.05 were LoF in our screen, suggesting predictive models of splicing will be valuable for prioritizing intronic variants for functional assessment. Whereas many LoF variants disrupt canonical splice sites, LoF variants were also observed in splice regions (3-8 bp from an exon), deeper in introns, in the 5’ UTR, and upstream of the transcriptional start site (TSS) (**Figure 4E**). Most variants in these regions lack definitive interpretations in ClinVar.

One such variant is c.885-3C>G, a VUS upstream of exon 11 predicted by SpliceAI to potentially cause acceptor site loss (SpliceAI score = 0.73). This variant scored comparably to known pathogenic variants adjacent to exon 11 in our screen (**Figure 4E**, middle). One notable variant deep within intron 15, c.1732-264A>T, is deemed “likely pathogenic” in ClinVar and scores intermediately by SpliceAI (0.39). This variant also scored as LoF in our assay (function score = 4.96; **Figure 4E**, right), corroborating its pathogenicity. In the *MLH1* 5’-untranslated region (5’- UTR), we scored c.-7_1del, an 8-bp deletion that disrupts the initiation codon as LoF, as well as 3.6% of other variants assayed in the 5’-UTR or upstream of the TSS.

To validate LoF effects of specific variants, we performed amplicon sequencing of edited regions, using the same gDNA from HAP1:PEmax cells from which pegRNAs were sequenced. Indeed, LoF variants including c.885-3C>G, c.1732-264A>T, and c.-7_1del were strongly enriched in their respective regions post-6TG selection, corroborating pegRNA-based function scores (**Figures S10C and S10D**). The lack of enrichment seen for nearly all SNVs in the 5’UTR and upstream region was consistent with the low function scores of these variants. Notably, one variant in close proximity to the TSS, c.-219G>T, was deemed LoF in the screen but not strongly enriched in ET sequencing, indicating that its pegRNA-derived function score may be conflated by CRISPR inhibition (CRISPRi) or some other unintended effect. While this example illustrates the importance of validating hits, this experiment nevertheless shows that our PE platform can be used to identify LoF variants acting via diverse mechanisms across large non-coding regions.

## DISCUSSION

Here, we demonstrate a PE platform that enables scalable functional interrogation of coding and non-coding variants in haploid human cells. We optimize key components of this platform for more efficient editing, demonstrate negative selection against LoF variants in an essential gene, and score a total of 763 variants in *MLH1*, newly identifying LoF variants in both coding and non-coding regions. Further, we extensively characterize the relationship between editing activity and the ability to accurately score variants.

In the context of well-established MAVEs for assessing variants by genome editing, such as base editing screens^18,19^, saturation genome editing via HDR^17^, and saturation prime editing^24^, our platform offers the advantage of being able to install any set of short variants virtually anywhere in the genome, provided an active pegRNA can be designed.

Current limitations stem from the fact that only a limited fraction of pegRNAs lead to robust editing. This explains the incomplete scoring of variants for which we designed pegRNAs. As we show for variants in both *SMARCB1* and *MLH1*, separation of signal from noise depends strongly on discerning more active pegRNAs, as editing percentages vary highly. In our platform, this is made possible via inclusion of STs with each pegRNA assayed. Nevertheless, in these proof-of-concept experiments we observed higher false negative rates for LoF variants than demonstrated in recent MAVE implementations^16,45^. We show this effect can be mitigated via more stringent pegRNA filtering, though this comes at a cost of reduced coverage.

Many variants throughout the *MLH1* gene had mildly positive function scores but did not pass the FDR cutoff (*q* < 0.01), and therefore were not deemed LoF. It remains to be determined whether such variants may be hypomorphic alleles. Alternatively, low endogenous editing rates may limit the strength of positive selection measurable via sequencing of pegRNAs alone. In the future, scoring multiple highly active pegRNAs per variant will improve precision.

Importantly, as PE reagents continue to improve and pegRNA design tools mature, we anticipate a larger fraction of pegRNAs in future experiments will produce accurate data. Based on this work, specific recommendations for performing future PE screens in HAP1 include: 1.) allowing ample time for edits to accrue, 2.) using a stabilized pegRNA scaffold, such as F+E v1, and 3.) including STs to filter out inactive pegRNAs. Though we saw a consistent benefit of ouabain co-selection, increases in editing rates were modest, suggesting this optimization may not always be necessary depending on the application. Implementing recently described pegRNA scoring tools^28,29^ in experimental design promises to improve coverage and reduce the number of pegRNAs with low editing activities.

While the strong selection for LoF variants afforded by 6TG proved valuable for scoring *MLH1* variants accurately, ultimately, additional assays will enable variants to be studied in more genomic regions. By establishing requirements for identifying LoF variants via negative selection, we illustrate a path forward for screening variants in over 2,000 genes that are essential in HAP1^41^. We envision this may be particularly valuable for prioritizing sets of variants identified in clinical sequencing for further study, and for testing effects of variants predicted to act via specific mechanisms across a large number of genes.

For assays of variants observed in patients, there remains a need for rigorous clinical benchmarking prior to using assay results to aid variant classification. With more data, there will be opportunities to refine function scores to better reflect variant-level confidence, incorporating both strength of pegRNA selection and ST editing rate. While we showed one strategy for validating variant effects by endogenous locus sequencing, in the future, improved pegRNA design will likely allow multiple, orthogonally designed pegRNAs per variant to be tested, either together or in successive experiments to validate hits.

We predict that implementing key improvements we have outlined will ultimately make data from genome-wide PE screens valuable for variant classification, as has been established for other MAVEs. Further, scaling this framework to test computationally prioritized sets of non-coding variants may yield data sets suited for training and refining predictive models. By presenting a means of engineering and assaying defined human variants across large sequence spaces, our platform for pooled PE promises to substantially improve identification of variants underlying human disease.

## SUPPLEMENTARY FIGURES

**Supplementary Figure 1.**
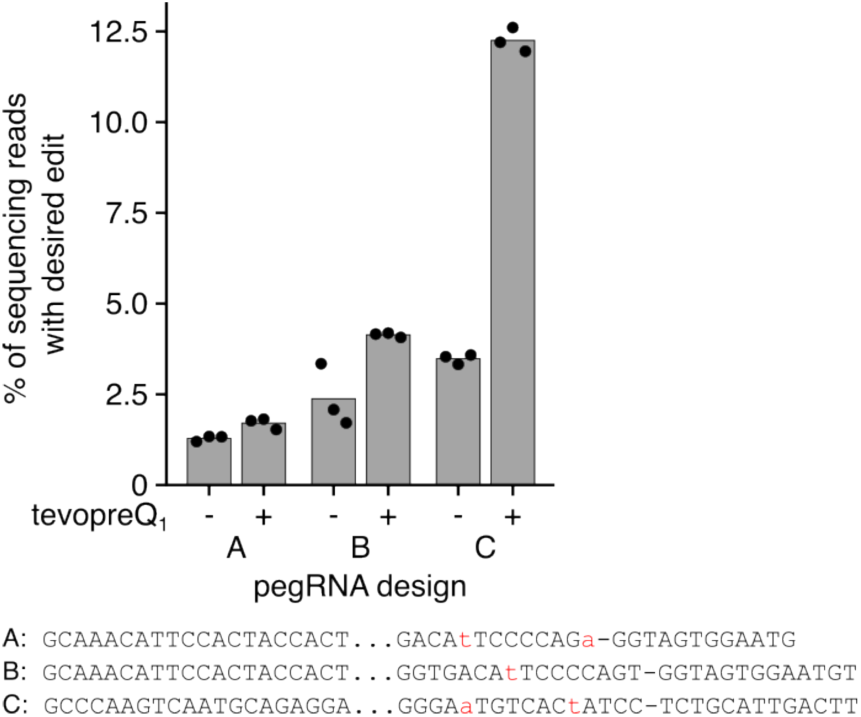
pegRNA optimization for *ATP1A1*-T804N edit in HEK293T. Comparison of PE efficiencies in HEK293T cells for installation of the T804N edit in *ATP1A1* using three pegRNA designs with and without the tevopreQ_1_ motif. Values correspond to percentages of correct editing 4 days after transfection of pegRNA and PEmax-MLH1dn plasmids as determined by CRISPResso2^46^ analysis of NGS reads. Individual transfection replicates are shown as dots. Below are sequences of pegRNA designs in the format “spacer…RTT-PBS”, with programmed variants highlighted in red.

**Supplementary Figure 2.**
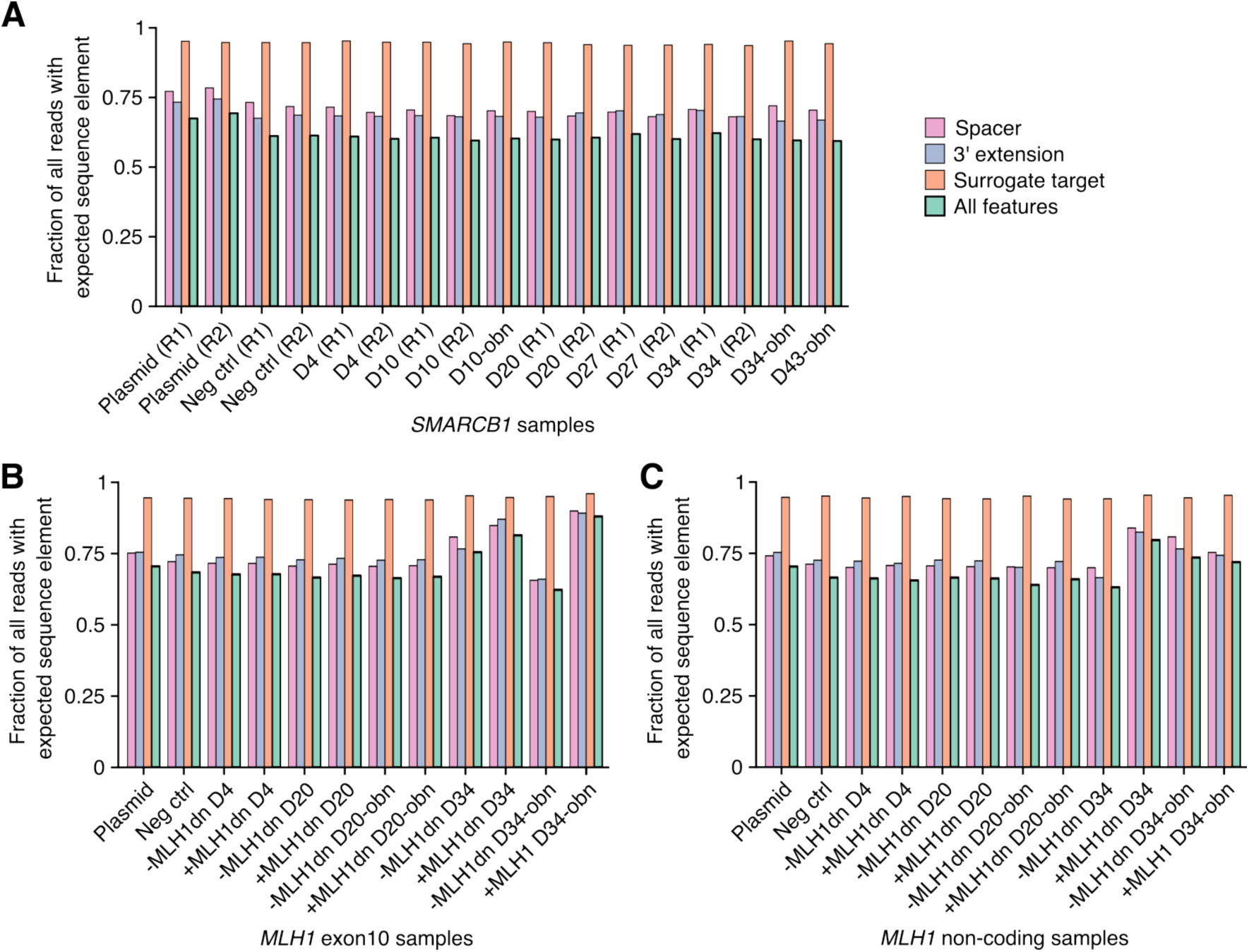
NGS read pre-processing of pegRNA-ST cassettes across experiments. Results from pre-processing pegRNA-ST cassette sequencing reads for each sample across experiments: *SMARCB1* (**A**), *MLH1* exon 10 (**B**), and *MLH1* non-coding (**C**). Values correspond to the fraction of reads containing each correct sequence element (spacer, 3’ extension, surrogate target or all elements) expected in the pegRNA construct, as identified by the read’s pegRNA-specific barcode. Spacer and surrogate target sequences were allowed to differ by up to two base substitutions from the expected sequence to account for sequencing errors.

**Supplementary Figure 3.**
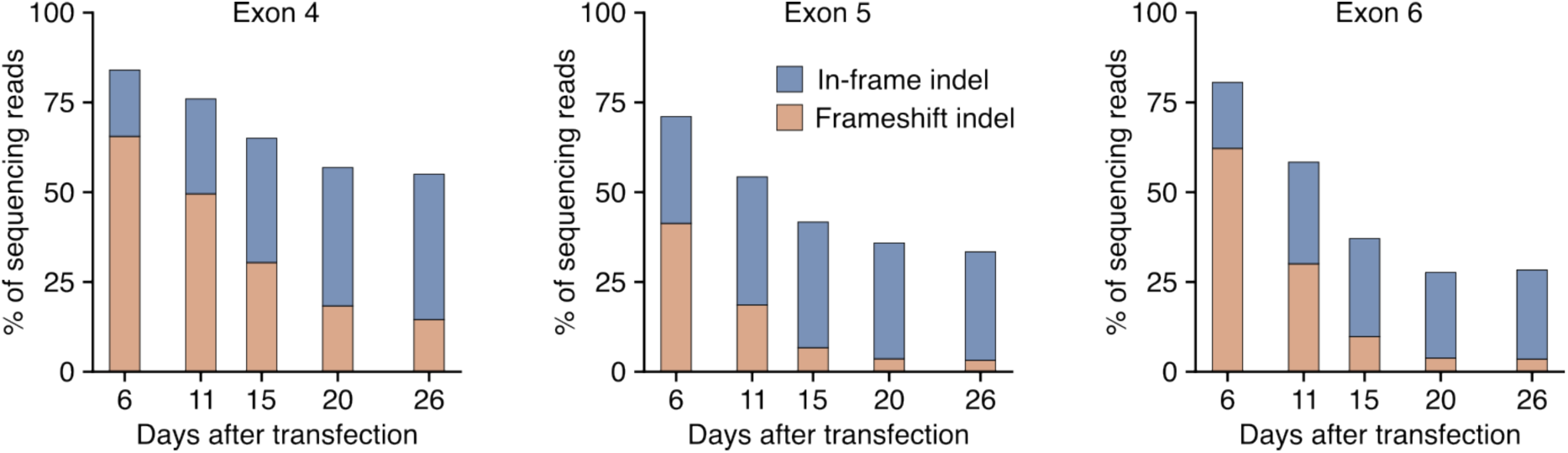
*SMARCB1* essentiality in HAP1. A time course was performed to assess in-frame and frameshifting indel percentages in exons 4, 5, and 6 of *SMARCB1* after Cas9-mediated editing in HAP1 cells. Indel rates were quantified from NGS reads using CRISPResso2.

**Supplementary Figure 4.**
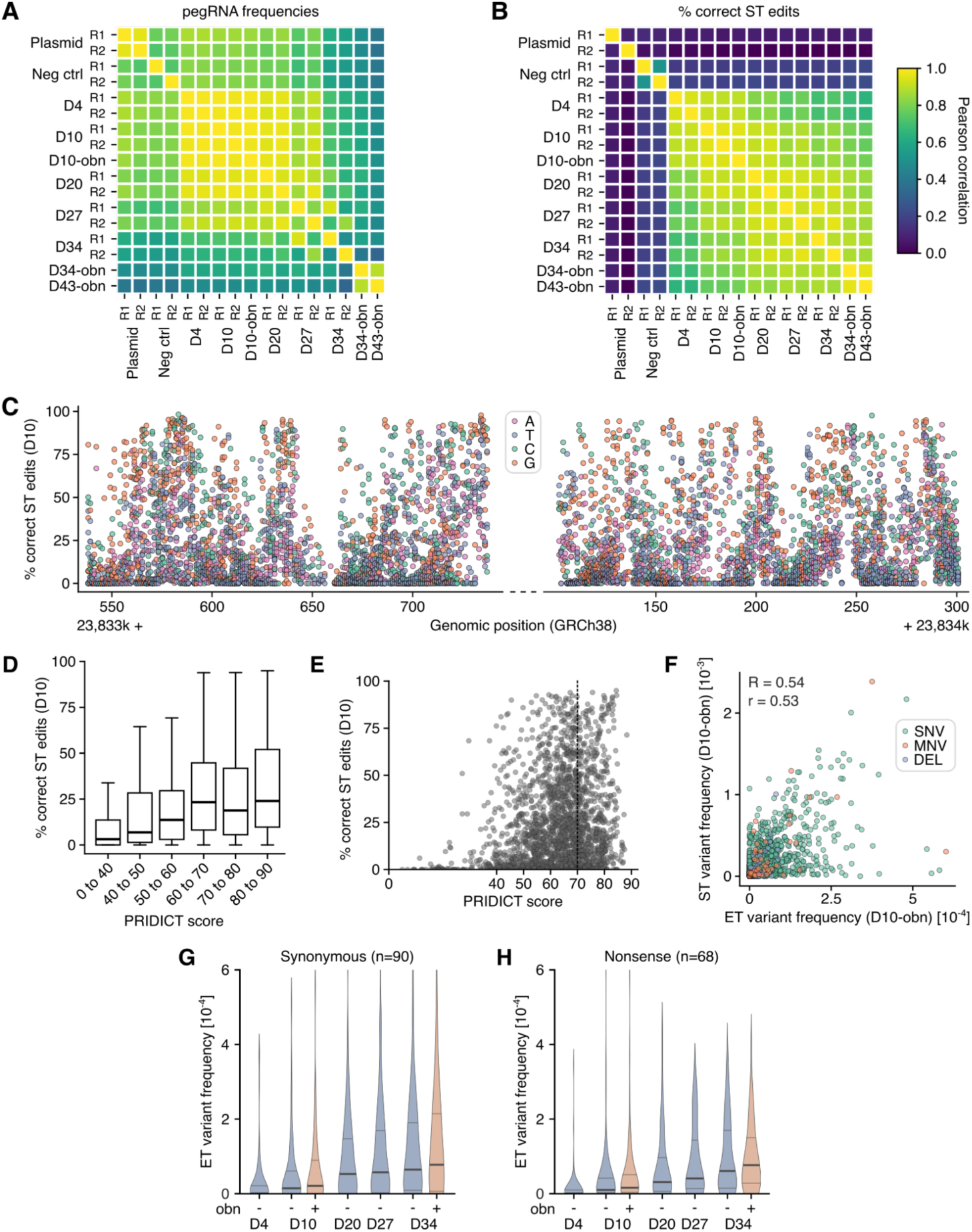
Characterization of pooled *SMARCB1* variant installation. (**A** and **B**) Heatmap of pairwise Pearson correlation coefficients between pegRNA frequencies (**A**) and correct ST editing percentages (**B**) across all collected samples of the *SMARCB1* variant screen, including the pegRNA plasmid pool (Plasmid) and transduced HAP1 wildtype cells (Neg ctrl). (**C**) Scatter plot of correct ST editing percentages 10 days post-transduction for each pegRNA (*n* = 6,902) targeting *SMARCB1* regions, colored by nucleotide substitution.(**D** and **E**) Boxplot and scatter plot showing correct ST editing percentages on day 10 as a function of pegRNA PRIDICT scores. In (**D**), bold line, median; boxes, IQR; whiskers extend to points within 1.5x IQR. In (**E**), the dashed vertical line marks the recommended PRIDICT threshold of 70. (F) The correlation between ST editing and ET editing for each variant is plotted. Variants are colored by edit type and Spearman (R) and Pearson (r) correlation coefficients are shown. (**G** and **H**) Variant frequencies at ETs across samples are plotted for all synonymous (**G**) and nonsense (**H**) variants. Frequencies were background corrected using NGS data from negative control samples. Similar enrichment of variants is observed over time for both synonymous and nonsense variants.

**Supplementary Figure 5.**
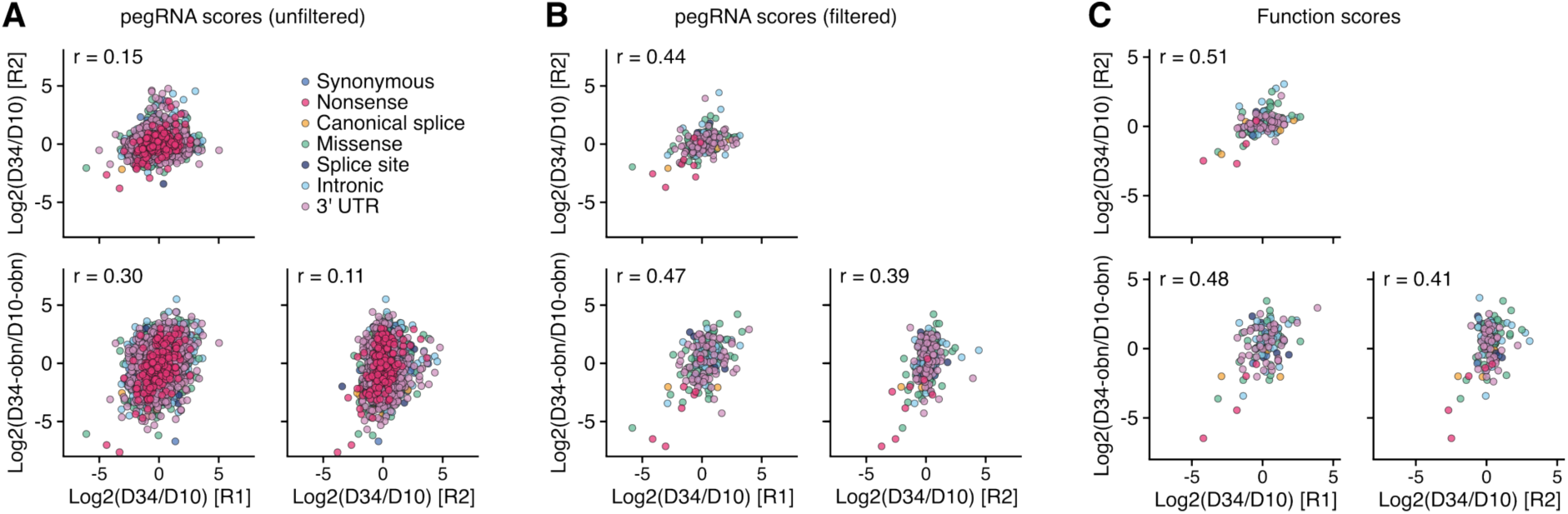
pegRNA score and function score correlations across *SMARCB1* experiments. Correlation plots of unfiltered (**A**) and filtered (**B**) pegRNA scores and function scores (**C**) for the *SMARCB1* variant screen between experiments with and without ouabain co-selection. Scores were calculated for pegRNA depletion between D10 and D34. pegRNAs passing filters were those with frequencies greater than 6×10^-5^ in D10 samples and correct ST editing percentages greater than 75%. Pearson correlation coefficients (r) are shown for each pairwise comparison.

**Supplementary Figure 6.**
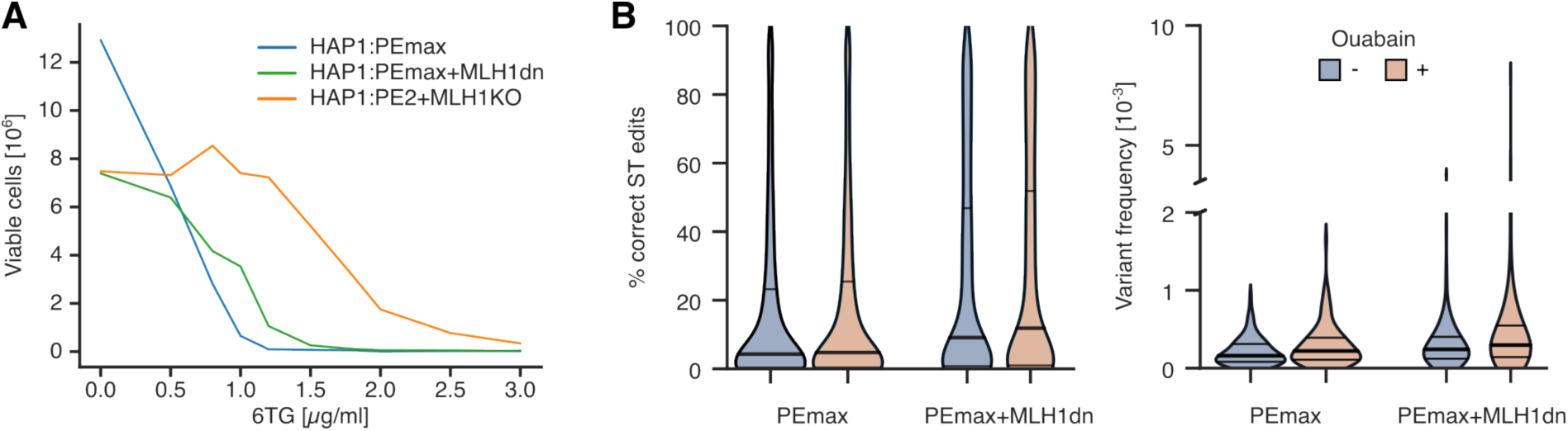
Effects of 6TG on growth and editing rates compared across HAP1 lines with modified MMR function. (A) Viable cell counts are plotted by 6TG dose following 6 days treatment for HAP1:PEmax, HAP1:PEmax+MLH1dn, and HAP1:PE2+MLH1KO. (B) Distributions of correct ST editing percentages for pegRNAs observed at frequencies greater than 1.4×10^-4^ in D20 samples (left), and distributions of ET variant frequencies in D20 samples (right) are shown for each condition. ET variants for which the log2-ratio of D20 frequency over D4 frequency was below 1.0 were excluded to limit potential impacts of sequencing error.

**Supplementary Figure 7.**
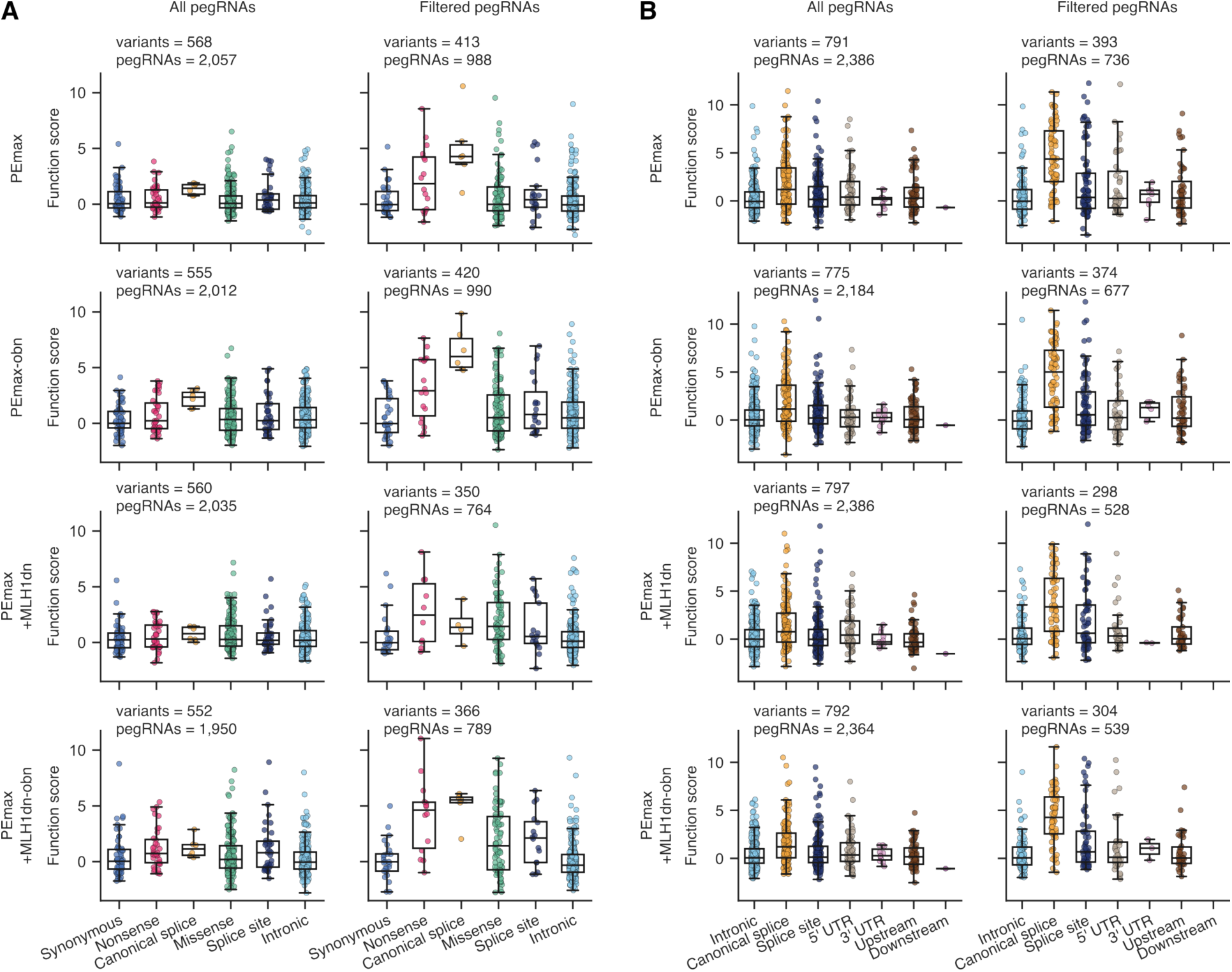
Filtering pegRNAs based on ST editing enables identification of LoF variants. For each *MLH1* experiment, function scores were calculated for each variant assayed, using either all pegRNAs observed above a frequency threshold, or only the subset of those with correct ST editing percentages above a set threshold. Function scores, grouped by variant consequence, are shown before and after pegRNA filtering for the *MLH1* exon 10 screen (**A**) and the *MLH1* non-coding screen (**B**). ST editing thresholds to filter pegRNAs were set to 5% for HAP1:PEmax cells and 25% for HAP1:PEmax+MLH1dn cells.

**Supplementary Figure 8.**
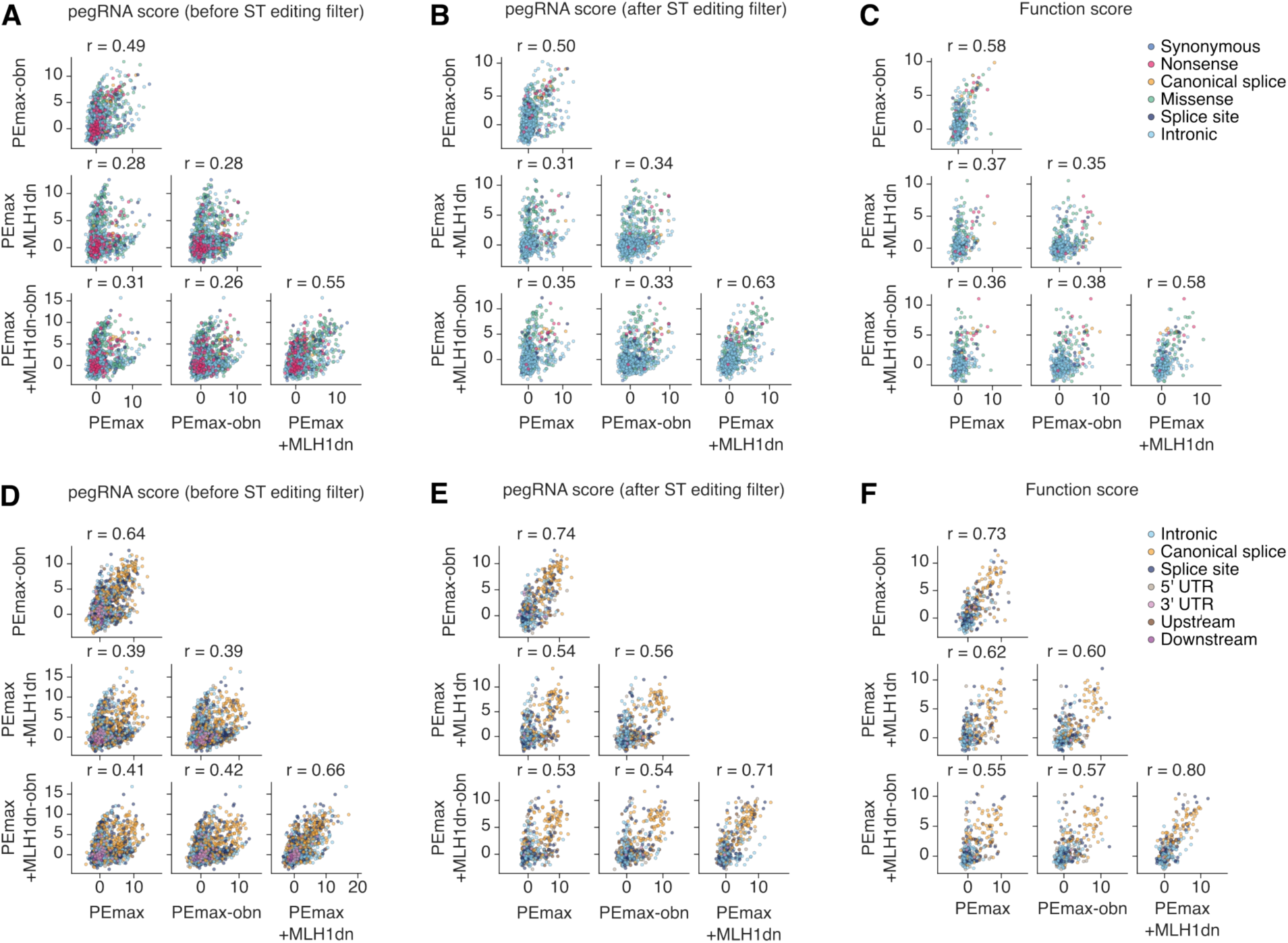
pegRNA score and function score reproducibility across experiments. Correlations of unfiltered and filtered pegRNA scores and function scores across conditions for the *MLH1* exon 10 screen (**A**-**C**) and the non-coding screen (**D**-**F**) are shown. Correct ST editing thresholds were set to 5% for HAP1:PEmax cells and 25% for HAP1:PEmax+MLH1dn cells to remove inactive pegRNAs and produce function scores from the filtered set. Pearson correlation coefficients (r) are shown for each comparison.

**Supplementary Figure 9.**
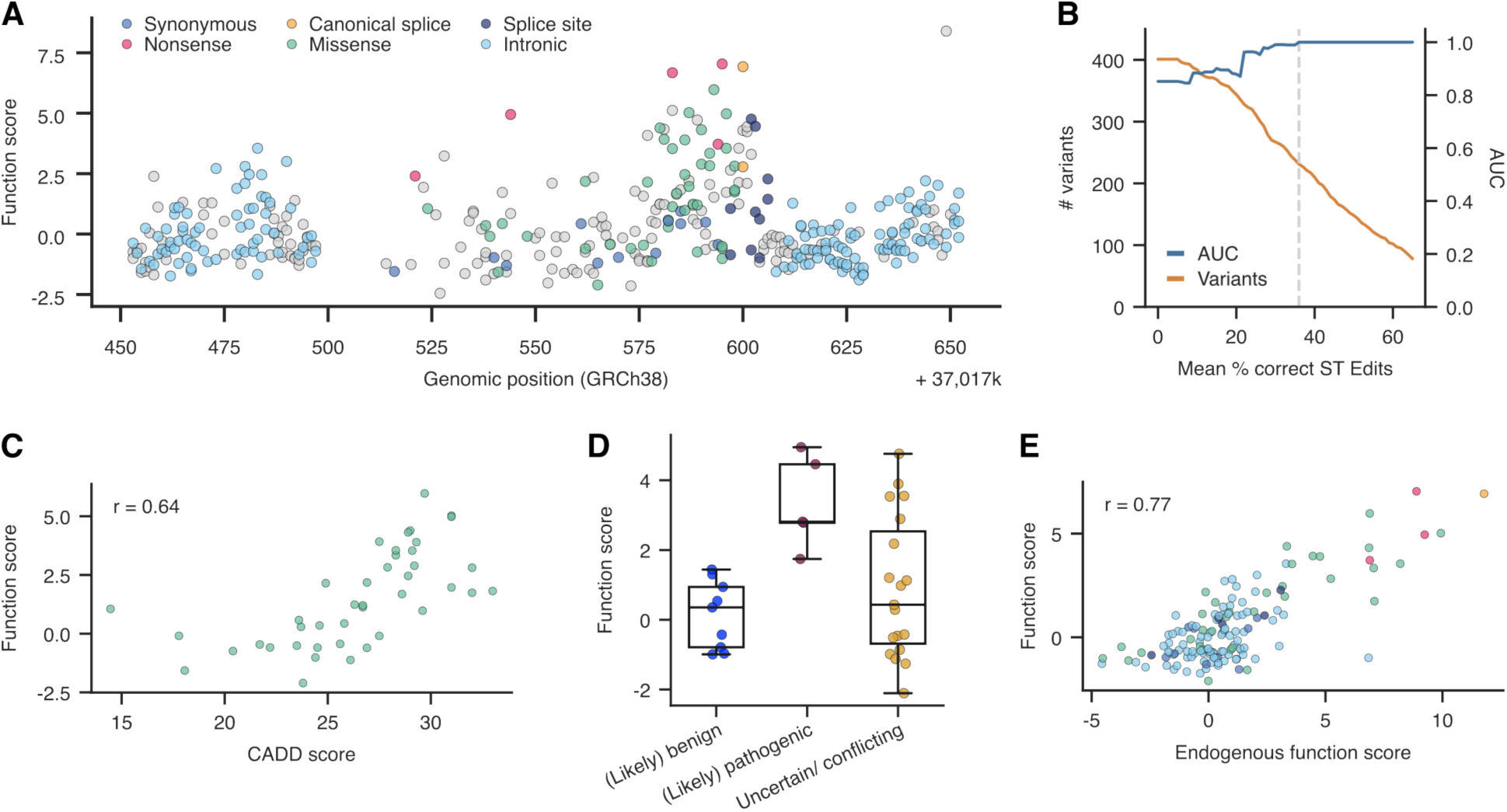
More stringent filtering of pegRNAs further improves data quality for variants assayed in *MLH1* exon 10. (A) Function scores for *n* = 231 variants scored using an average correct ST editing threshold of 36% are plotted by position. (Gray dots indicate variants scored in Figure 3C with ST editing percentages below 36%.) (B) AUC values for distinguishing LoF variants (blue) are plotted as a function of the mean ST editing threshold applied. For this analysis, synonymous variants were defined as neutral and nonsense and canonical splice variants as LoF. The orange line indicates the number of variants retained at each threshold, and the dashed line indicates the high-stringency threshold of 36%, above which AUC = 1.0. (C) The correlation between function scores and CADD scores is plotted for *n* = 42 missense variants passing the high-stringency threshold. (D) The boxplot shows function scores for *n* = 33 variants passing the high-stringency threshold by ClinVar pathogenicity status (bold line, median; boxes, IQR; whiskers extend to points within 1.5x IQR). (E) The correlation between function scores (pegRNA-derived) and endogenous function scores is plotted for *n* = 141 variants passing the high-stringency threshold.

**Supplementary Figure 10.**
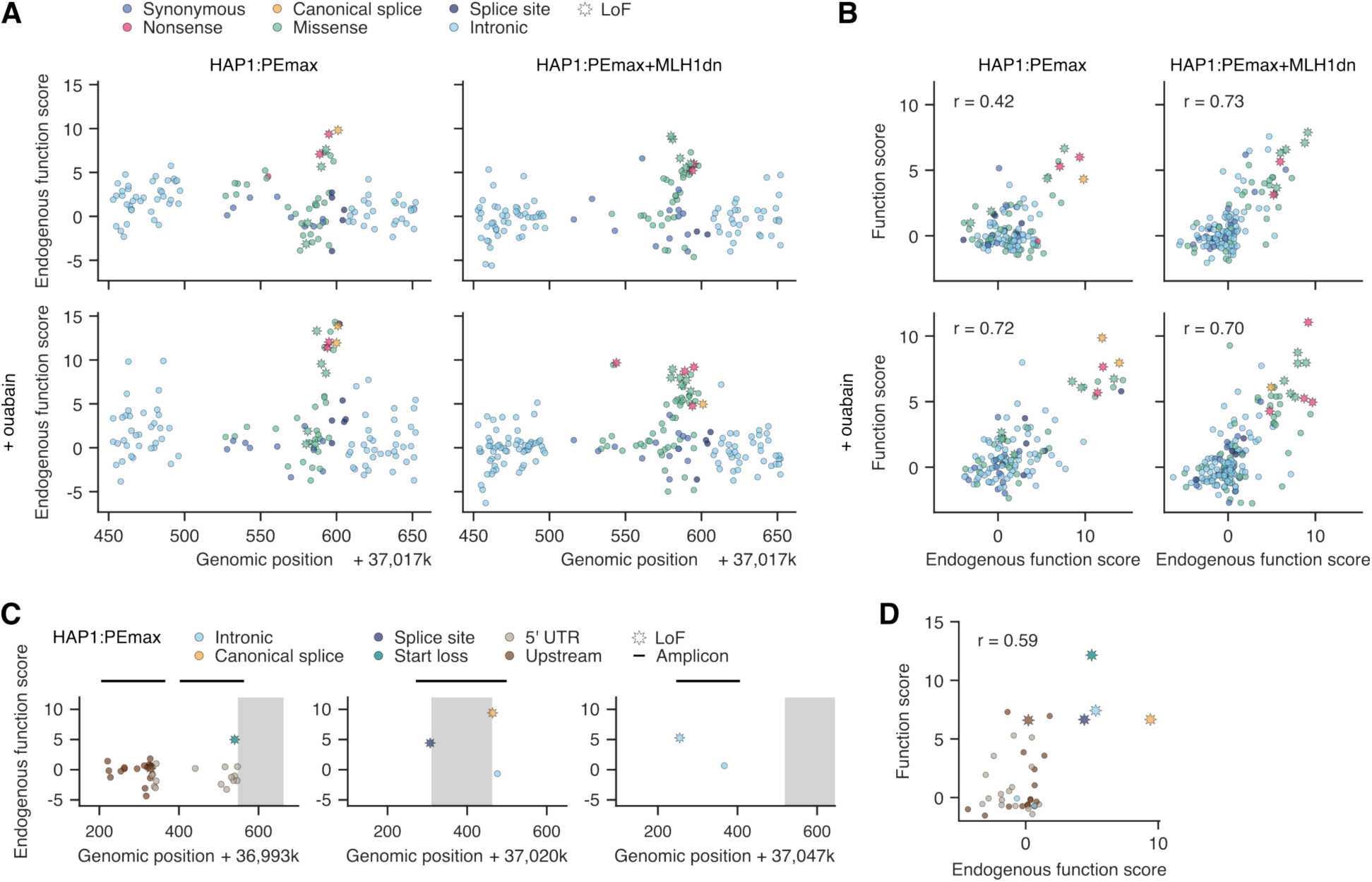
Amplicon sequencing of edited *MLH1* loci validates functional effects of variants installed via PE. (A) Endogenous function scores of exon 10 variants are plotted by genomic position across conditions. Variants deemed LoF via pegRNA sequencing of all conditions are starred (B) Correlations between function scores and endogenous function scores are plotted for each experiment (C) Endogenous function scores are plotted by genomic position for each non-coding region validated. Endogenous function scores were derived by sequencing edited loci in HAP1:PEmax cells pre- and post- 6TG selection (D) Correlation between function scores and endogenous function scores from HAP1:PEmax for regions in (**C**).

**Supplementary Figure 11.**
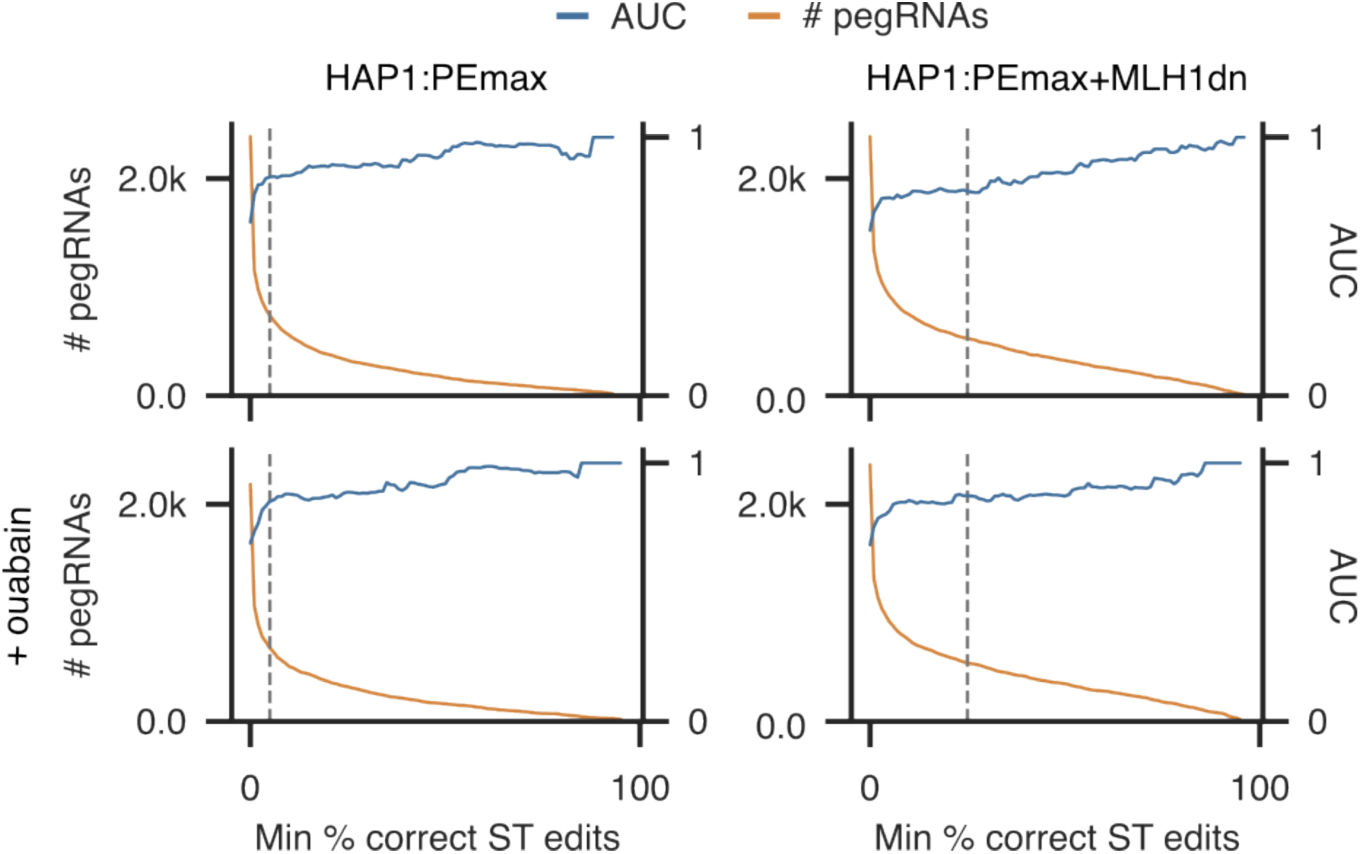
Accurate identification of LoF variants in the *MLH1* non-coding screen is facilitated by implementing ST editing thresholds. AUC measurements for distinguishing pLoF variants (blue line) are plotted as a function of ST editing threshold, shown for each experimental condition. The number of pegRNAs retained at each ST editing threshold is plotted in orange. For these experiments, pNeut variants were defined as intronic variants greater than 8 bp from exonic sequence and pLoF variants were defined as canonical splice site variants.

## METHODS

### Plasmids

For transient co-expression of PEmax and MLH1dn we used the pCMV-PEmax-P2A-hMLH1dn plasmid (*Addgene #174828*). The pU6_pegRNA-GG-Acceptor plasmid (*Addgene #132777*) was modified to exchange the mRFP1 cassette with a BsmBI cloning cassette, allowing for straightforward insertion of pegRNAs. Gibson assembly reactions to create these plasmids were performed using NEBuilder HiFi DNA Assembly Master Mix (New England Biolabs, #E2621L).

For stable integration of PEmax with and without MLH1dn, a lentiviral vector was prepared by PCR amplification of the corresponding cassettes from pCMV-PEmax-P2A-hMLH1dn and subsequent insertion into pLenti-PE2-BSD (*Addgene #161514*), which had been digested with EcoRI and XbaI (#R0145S & #R3101S, *NEB*). Single pegRNAs were stably integrated using lentiviral vectors cloned by assembling pegRNA oligo duplexes into BsmBI-digested (#R0739L, *NEB*) Lenti_gRNA-Puro (*Addgene #84752*). The dual-pegRNA lentiviral vector, used for pegRNA library cloning, was prepared by amplification of a gene fragment encoding the mU6 promoter and a BsmBI cloning site (*Twist Bioscience*) followed by assembly into a KflI- and Eco72I- digested (#FD2164 & #FD0364, *Thermo Scientific*) Lenti_gRNA-Puro plasmid with a pre-inserted co-selection pegRNA under the hU6 promoter. For bacterial amplification of pegRNA scaffolds, oligo duplexes, encoding the different scaffold designs (original, F+E, F+E v1, F+E v2) and flanking BsmBI recognition sites, were inserted into the KpnI-digested (#R3142S, *NEB*) pU6_pegRNA-GG-Acceptor plasmid via Gibson assembly.

All-in-one CRISPR knock-out constructs targeting *MLH1* exon 10 and *SMARCB1* exons 4, 5, and 6 were generated from pSpCas9(BB)-2A-Puro (PX459 V2.0, *Addgene #62988*) via insertion of spacer sequences into the BbsI cloning site (#R3539L, *NEB*). **Supplementary Table 9** contains sequences for all oligos used in this study.

### Cell culture

HEK293T cells were cultured in Dulbecco’s Modified Eagle Medium (DMEM) (#10564011, *Gibco*) supplemented with 10% fetal bovine serum (FBS) (#A5256701, *Gibco*) and 1% Penicillin-Streptomycin (Pen-Strep) (#15140122, *Gibco*). Cells were cultured at 37 °C with 5% CO_2_ and passaged using Versene solution (#15040066, *Thermo Fisher*). HAP1 cells were cultured in Iscove’s Modified Dulbecco’s Medium (IMDM) (#12440053, *Gibco*) supplemented with 10% FBS, 1% Pen-Strep, and 2.5 µM 10-deacetyl-baccatin-III (DAB) (#S2409-SEL, *Stratech*) at 37 °C with 5% CO_2_. Cells were passaged every 2-3 days using 0.25% trypsin-EDTA (#25200056, Gibco) to keep them below 80% confluency. For culture of HAP1:PEmax cell lines, media was additionally supplemented with 5 µg/mL blasticidin S (R21001, *Gibco*). All cell lines were confirmed to be free of mycoplasma.

### *SMARCB1* indel depletion in HAP1

Approximately 4×10^6^ HAP1 cells were transfected using Xfect Transfection Reagent (#631318, *Takara Bio*) for each pSpCas9(BB)-2A-Puro construct. 24h after transfection, the culture media was exchanged and supplemented with 1 µg/mL puromycin (#13884, *Cayman Chemical*). Approximately 10^7^ cells were harvested on days 6, 11, 15, 20, and 26 after transfection and stored at −80 °C. Cell pellets were processed for extraction and purification of genomic DNA using DNeasy Blood and Tissue Kit (#69506, *Qiagen*). Target loci were amplified from purified gDNA by PCR using NEBNext Ultra II Q5 Master Mix (#M0544X, *NEB*). Adapters for dual-indexed Illumina sequencing were attached in a second PCR step. Sequencing reads were processed using the Cas9 mode of CRISPResso2 and its default parameters^46^.

### pegRNA library design

All pegRNA libraries were designed using a Jupyter Notebook implementation of the open-source python package PEGG^30^. For *SMARCB1* and *MLH1* saturation mutagenesis libraries, all possible SNVs in a 200 bp window centered on each target exon were created. We additionally introduced nonsense mutations at each codon via multinucleotide substitutions. For *SMARCB1* saturation mutagenesis libraries, we also included single-codon deletions. MNVs for the experiment comparing editing rates of SNVs and MNVs were designed from a random subset of the SNVs programmed as part of the *SMARCB1* saturation mutagenesis experiment. Up to two additional synonymous mutations were programmed in the four neighboring codons nearest the target codon. ClinVar variants within *MLH1* were accessed on 11/29/2022, filtering for short variants (less than 10 bp) with variant start positions mapping to non-coding regions.

*SMARCB1*-targeting libraries were designed to include up to 9 pegRNAs per variant, while *MLH1* libraries contained up to 12 pegRNAs per variant. To diversify the set of pegRNAs per variant, we enforced the use of distinct spacer sequences (3 distinct spacers for *SMARCB1* pegRNAs and 2 distinct spacers for *MLH1* pegRNAs, respectively) and selected the 3 top-scoring pegRNAs within each set of pegRNA designs using a common spacer. All pegRNAs were appended with the tevopreQ_1_ motif followed by a T_7_ termination sequence. Additionally, each pegRNA was coupled to a 55-nt surrogate target (ST) sequence (replicating the endogenous target) and a unique 16-nt pegRNA barcode. The scaffold sequence was replaced with a BsmBI cloning cassette for ordering oligos, and pegRNA oligos were divided into multiple sub-libraries, each with a unique 10-nt library barcode at the oligo’s 3’ end to allow specific amplification of individual libraries from the oligo pool. To create oligos of equal size (243 nt), a stuffer sequence was inserted between the T_7_ termination sequence and ST sequence. Finally, designed pegRNA oligos containing BsmBI recognition sites that would interfere with cloning were discarded. pegRNA oligo libraries were ordered and synthesized as a custom oligo pool (*Twist Bioscience*).

### pegRNA library cloning

pegRNA libraries were cloned from oligo pools into lentiviral vectors in a two-step procedure. pegRNAs were amplified from the oligo pool via PCR using KAPA HiFi HotStart ReadyMix (#KK2602, *Roche*). pegRNA oligo subsets were specifically amplified from the oligo pool using primers specific for the library BC (**Supplementary Table 9**). Thermocycling was performed according to guidelines for KAPA HiFi HotStart ReadyMix except for elongated extension steps of 2 min. PCR products were purified and concentrated using AMPure XP SPRI Reagent (#A63881, *Beckman Coulter*) and subsequently by agarose gel electrophoresis to isolate 260 bp amplicons.

The dual pegRNA lentiviral vectors with pre-integrated co-selection pegRNAs were prepared for pegRNA library cloning by BsmBI-digestion and Quick CIP (#M0525L, *NEB*) treatment followed by gel purification. pegRNA oligo amplicons were assembled with the cut lentiviral vector via NEBuilder HiFi DNA Assembly Master Mix. The assembled plasmid pool was purified and concentrated with AMPure XP SPRI Reagent and used for transformation of Electrocompetent Endura Cells (#60242-2, *Biosearch Technologies*) following manufacturer’s instructions and a previously published protocol^47^. The plasmid pool was extracted and purified using ZymoPURE II Plasmid Maxiprep Kit (#D4202, *Zymo Research*).

A second cloning step was next required to insert a pegRNA scaffold sequence. Therefore, the plasmid pool was subjected to BsmBI-digestion, Quick CIP treatment, and gel-electrophoretic purification. Separately, pegRNA scaffolds were prepared for cloning by BsmBI-digestion of pScaffold plasmids followed by gel purification. The digested vector library and the pegRNA scaffold were ligated using T4 DNA Ligase (#M0202, *NEB*). SPRI-purified and concentrated ligation product was used for transformation of Electrocompetent Endura Cells, as before, and plasmid pools were purified for lentivirus production. A 1,000-fold coverage of pegRNA library size was ensured at each transformation step, with the exception of the *MLH1* non-coding library, where the minimum coverage exceeded 250-fold at each step.

For cloning of the pegRNA pool to compare performance of different scaffolds, the first step of pegRNA library cloning remained the same. Scaffolds for all tested designs were prepared from corresponding pScaffold plasmids via BsmBI-digest and pooled at equimolar ratios. This scaffold mix was used in the subsequent ligation reaction with the BsmBI-digested plasmid pool before proceeding as detailed above.

The pegRNA pool for the *SMARCB1* variant screen was cloned into two lentiviral vectors with distinct co-selection pegRNAs. One vector encoded the *ATP1A1*-T804N pegRNA for ouabain co-selection, while the other vector contained a *HPRT1*-A161E pegRNA. Lentiviral particles were produced from both plasmid pools, which were subsequently used for transductions in *SMARCB1* variant screening experiments. Although intended to provide an alternative possible co-selection strategy, the pegRNA pool in the *HPRT1*-A161E co-selection vector served only as a duplicate library for assaying variants without co-selection.

### Lentivirus production and titering

For lentivirus production, each transfer plasmid was mixed with packaging and envelope plasmids (pLP1, pLP2, VSV-G) to 33 µg total DNA which was used for transfection of 2×10^7^ HEK293T cells in a 15-cm culture dish with Lipofectamine 2000 (#11668019, *Invitrogen*). Viral supernatants were collected 2 days and 3 days post-transfection, before pooling and concentrating with PEG-8,000 (#V3011, *Promega*). Aliquots of concentrated lentivirus were stored at −80 °C until used. Aliquots of lentivirus particles were titered by ddPCR as described^48^.

### Generation of HAP1 PE cell lines

HAP1:PEmax and HAP1:PEmax+MLH1dn cell lines were generated via transduction of 3×10^6^ HAP1 cells at low MOI (less than 0.3) with lentiviral particles packaged using pLenti_PEmax-BSD transfer plasmids with and without MLH1dn expression. 2 days after transduction, the media was replaced with media containing 5 µg/mL blasticidin. Selection of transduced cells was performed for 14 days before expanding single clones. Final clones were chosen based on functional validation of PE activity via *ATP1A1* editing followed by ouabain selection. Complete genomic integration of the PEmax+MLH1dn sequence was validated via PCR of this cassette from genomic DNA followed by gel electrophoresis.

The HAP1:PE2+MLH1KO line was used only for titering 6TG dose. It was created by first transducing parental HAP1 cells with pLenti-PE2-BSD at low MOI (as for PEmax cell lines), then transfecting successfully transduced cells with a pX459 construct targeting *MLH1*. Individual clones were isolated and screened by Sanger sequencing, and a line with a frameshifting indel in exon 18 was selected.

### *ATP1A1*-T804N pegRNA optimisation

Around 10^5^ HEK293T cells were co-transfected with pCMV-PEmax-P2A-hMLH1dn and pU6_pegRNA plasmids (6 different pegRNA designs) using Xfect Transfection Reagent. 4 days after transfection, cells were harvested and gDNA was extracted and purified. The target locus was amplified from gDNA by PCR using NEBNext Ultra II Q5 Master Mix (#M0544L, *NEB*). Adapters for dual-indexed Illumina sequencing were attached in a second PCR step. Sequencing reads were processed using the prime editing mode of CRISPResso2 to determine the fraction of reads with desired edits.

### *ATP1A1*-T804N enrichment with ouabain

Approximately 10^7^ HAP1:PEmax+MLH1dn cells were transduced at low MOI (approximately 0.1) with lentivirus particles produced from a Lenti_pegRNA-Puro plasmid encoding the ATP1A1-T804N pegRNA under the hU6 promoter. Cell selection was initiated 1 day later by supplementation of the culture media with 1 µg/mL puromycin and was maintained for 3 days. On day 4 post-transduction, the cell pool was split in two for continued culture with and without ouabain treatment. Selection of cells with the *ATP1A1*-T804N edit was started by addition of 5 µM ouabain (#O3125, *Sigma-Aldrich*) and maintained until the end of the experiment. Cells were harvested for gDNA extraction and purification on days 4 and 11 post-transduction. The target locus was PCR amplified and sequenced by NGS. The fraction of sequencing reads with perfect and partial editing were determined via counting matches for corresponding 18-nt subsequences and dividing by total number of sequencing reads.

### 6TG dose titration

1.2x10^7^ HAP1:PEmax, HAP1:PEmax+MLH1dn, and HAP1:PE2+MLH1KO cells were treated for 6 days with a range of 6TG (#A4882, *Sigma-Aldrich*) concentrations: 0µM, 0.5µM, 0.8µM, 1µM, 1,2µM, 1.5µM, 1.8µM, 2µM, 2.5µM, 3µM, 3.5µM, 4µM. Viable cells post drug challenge were counted using the Vi-Cell analyzer (Beckman).

### Pooled PE screening

For all pooled PE screens, prepared pegRNA lentivirus aliquots were thawed on ice and used to transduce HAP1:PEmax lines at low MOIs (0.1-0.5), maintaining an average pegRNA coverage of at least 1,000-fold for each library. As a negative control, we also transduced HAP1 cells not expressing PEmax at a higher MOI of approximately 10, achieving 100x pegRNA coverage. One day after transduction, the culture media was exchanged and supplemented with 1 µg/mL puromycin for 3 days of selection, at which point MOIs were confirmed by examining cell confluency. Aliquots of cells were harvested periodically throughout the experiment, ensuring at least 1,000-fold average coverage of the library at each timepoint. Negative control samples were harvested at the earliest timepoint (D4 or D5) and served to determine background variant rates in STs. Additional experimental details for each pooled PE experiment are as follows:

### pegRNA scaffold activity screen

Approximately 4×10^7^ HAP1:PEmax+MLH1dn cells were treated with lentivirus particles, achieving an MOI of approximately 0.1. The negative control sample (transduced HAP1 cells without PEmax expression) was harvested on D5, while the experimental pool was harvested on D7 to allow additional time for editing.

### SNV vs. MNV pegRNA activity screen

pegRNA library cloning, lentivirus production, and screening were carried out in duplicate. The pegRNA library was tested in HAP1:PEmax+MLH1dn cells as part of the larger *SMARCB1* variant screen and analyzed separately. Comparison of SNV and MNV ST editing rates was performed using non-ouabain-treated samples harvested on D10, whereas the negative control sample was processed on D4.

### *SMARCB1* saturation mutagenesis screen

*SMARCB1* pegRNA library cloning, lentivirus production, and transduction of HAP1:PEmax+MLH1dn cells were performed in duplicate (once cloned into a vector co-expressing the *HPRT1*-A161E pegRNA and once cloned into a vector co-expressing the *ATP1A1*-T804N pegRNA for ouabain co-selection). To achieve at least 1,000-fold average coverage of the pegRNA library (*n* = 12,211 pegRNAs), total cell numbers were maintained above 1.3×10^7^ throughout the experiment. After completion of puromycin selection 4 days post-transduction, the cell pools were split into two. One pool was maintained in media containing 5 µM ouabain while the remaining two duplicates were left untreated. Cells were passaged every 2-3 days and pellets of at least 3×10^7^ cells were harvested on days 4, 10, 20, 27, and 34 post-transduction. The negative control sample was processed on D4.

### *MLH1* exon 10 and non-coding variant screens

The same experimental procedure was applied for the *MLH1* exon 10 saturation mutagenesis and non-coding ClinVar variant screens. Lentiviral aliquots of pegRNA pools were used for transduction of both HAP1:PEmax and HAP1:PEmax+MLH1dn cell lines (D0). To achieve at least 1,000-fold average coverage of the pegRNA library (*n* = 2,696 pegRNAs for the exon 10 pool and *n* = 3,748 pegRNAs for the non-coding ClinVar pool), at least 4×10^6^ cells were maintained throughout the experiment and harvested at each sampling. Puromycin selection was completed by D4 and on D13 the cell pools were split into two. One cell pool per cell line was treated with 5 µM ouabain while the other was left untreated, resulting in a total of four cell pools per experiment (HAP1:PEmax and HAP1:PEmax+MLH1dn each with and without ouabain co-selection). 20 days post-transduction, 6TG was added to the culture media at a concentration of 1.2 µg/ml for HAP1:PEmax and 1.6 µg/ml for HAP1:PEmax+MLH1dn. Cells were passaged every 2-3 days with addition of fresh selection media until D34. Cell pellets of at least 10^7^ cells were harvested on D4, D20 (pre-selection), and D34 (post-selection). The negative control sample was harvested on D4.

Cell pellets were processed for gDNA extraction and deep sequencing of both pegRNA-ST cassettes and ETs was performed. For *SMARCB1* and *MLH1* exon 10 saturation mutagenesis screens, both pegRNA-ST cassettes and ETs were sequenced, while for screens in which only pegRNA activity was assessed only the pegRNA-ST cassette was sequenced. For the *MLH1* non-coding variant screen the pegRNA-ST cassette was sequenced across all conditions. Additionally, the following amplicons were sequenced from HAP1:PEmax cells without co-selection to validate selective effects for a subset of variants: 164 bp spanning the transcription start site, 138 bp of the 5’UTR extending into exon 1, 197 bp including exon 11 and adjacent intronic sequence, and 133 bp within intron 15. The same regions were sequenced from control cells (unedited) to quantify baseline sequencing error.

### Extraction of genomic DNA and PCR amplification

Extraction of gDNA from cell pellets was performed using the DNeasy Blood and Tissue Kit (#69504, *Qiagen*) following the supplier’s protocol. For amplicon sequencing, genomic sites of interest were amplified by PCR using KAPA HiFi HotStart ReadyMix. Up to 2.5 µg gDNA was used as template per 100 µL PCR volume and reaction mixtures were supplemented with MgCl_2_ (#AM9530G, *Invitrogen*) added to a concentration of 5 mM. Reactions were supplemented with SYBR Safe (#S33102, *Invitrogen*) and run on a real-time PCR machine to prevent overcycling. Enough reactions were performed and subsequently pooled to maintain at least 1,000-fold average coverage of each pegRNA library when amplifying both pegRNA-ST cassettes and ETs. Primer annealing temperatures for all reactions were predetermined using gradient PCR prior to sample processing. pegRNA-ST amplicons were sequencing-ready after purification of the first PCR from gDNA. ET amplicons were pooled across independent reactions, SPRI-purified and used as template in subsequent PCRs to introduce Illumina sequencing adapters and sample indexes. For the *MLH1* exon 10 experiment, 1 additional PCR was performed for sample indexing, whereas for the *MLH1* non-coding experiment, an additional nested PCR to install sequencing adapters was performed prior to sample indexing.

### Illumina sequencing

Dual-indexed amplicons with Nextera or TruSeq adapters were sequenced using either an Illumina NextSeq 500 300-cycle kit or an Illumina NovaSeq 6000 SP 300-cycle kit. For deep sequencing of pegRNA-ST cassettes, the length of read 1 was set to 182 nt to capture the full pegRNA sequence and the length of read 2 was set to 118 nt to cover the ST, pegRNA BC and library BC sequences.

### Sequencing data analysis

All bcl files were demultiplexed and converted to fastq files using bcl2fastq2.

### ET analysis

Sequencing reads of ETs from pooled PE screens were processed using DiMSum (– vsearchMinQual 5, –maxSubstitutions 3, otherwise default parameters)^49^. Variant counts were further processed with custom Python scripts available on GitHub. Variants not programmed by pegRNAs within the pool were discarded.

*SMARCB1* ET variant frequencies were corrected for sequencing error by subtraction of background frequencies observed in the negative control sample (non-transduced wildtype cells), with any resulting negative value set to zero. A small number of variants were highly abundant in the negative control sample, likely owing to site-specific sequencing error. Therefore, variants with negative control frequencies above 4×10^-4^ were excluded from further analysis. Variant frequencies were averaged across duplicate samples where available.

For *MLH1* ET analyses, log2-ratios of variant frequency on D20 over D4 (saturation screen) or D20 over negative control (non-transduced wildtype cells) (non-coding screen) were calculated. Variants with log2-ratios below 1 were excluded from analysis (i.e., those not enriched over the course of editing). Endogenous function scores were calculated for variants that received a pegRNA-derived function score, as log2-ratios of variant frequency on D34 over D20 (post- and pre-6TG selection, respectively), normalized to the median score of synonymous variants (exon 10 screen) or intronic variants annotated as benign in ClinVar (non-coding screen). For the exon 10 screen, normalized endogenous function scores were averaged across conditions to yield a single endogenous function score per variant. These scores were once more normalized to the median synonymous score to produce a final endogenous function score per variant.

### pegRNA-ST read processing

A custom bash script was run on demultiplexed fastq files to extract and write new fastqs for each sequence element, including the protospacer, scaffold, 3’ extension, ST, pegRNA BC, and library BC using cutadapt (version 4.4). Using a custom Python script, sequence elements of each read were queried against a list of expected pegRNA-ST cassettes and reads with non-matching combinations of elements were discarded. For STs, only the 3’ end of each sequence was used for cross-checking such that reads with edited STs would not be discarded. Matches with up to 10% substitutions per element were allowed (to accommodate sequencing errors), however recombined pegRNA-ST cassettes were recognised and discarded in this step. For each sample, sequencing reads with identical pegRNA identities were tallied to determine pegRNA counts. Next, ST reads were grouped by pegRNA identity and queried for correct editing by searching for a string comprising the intended PE edits flanked by 5 bp on either side. Percentages of correct ST edits were calculated as the number of STs with correct editing over the total number of STs for each pegRNA with at least 10 sequencing reads. pegRNAs with greater than 5% correct ST editing in the negative control sample were excluded from downstream analyses.

### Calculation of pegRNA scores and function scores

For each sample, pegRNA counts were incremented by 1 and converted to frequencies. To measure the change in pegRNA frequency during selection, pegRNA scores were calculated as the log2-ratio of pegRNA frequency in the post-selection sample over pegRNA frequency in the pre-selection sample. To avoid assigning scores to poorly sampled pegRNAs, pegRNAs were filtered on pegRNA frequency in pre-selection samples. Where indicated, pegRNAs were also filtered for editing activity using the percentage of correct ST editing observed for each pegRNA.

For the *SMARCB1* variant screen, pegRNA filtering thresholds were set to 6×10^-5^ for pegRNA frequency and 75% correct ST editing for pegRNA activity. For *MLH1* variant screens, pegRNA frequency thresholds were set to 1.4×10^-4^ (exon 10 screen) or 1.0×10^-4^ (non-coding screen). Activity thresholds were set to 5% correct ST editing for all *MLH1* screens performed in HAP1:PEmax cells and to 25% correct ST editing for all screens performed in HAP1:PEmax+MLH1dn cells. PegRNA scores were normalized to the median pegRNA score of synonymous variants for the *SMARCB1* and *MLH1* saturation mutagenesis screens, or to the median pegRNA score of intronic variants for the *MLH1* non-coding screen.

Function scores for each variant were computed by averaging pegRNA scores for all pegRNAs programming the same variant. For the *SMARCB1* screen, this was achieved by averaging normalized pegRNA scores for each variant. For *MLH1* screens, unnormalized pegRNA scores from each experimental condition were used to determine condition-specific function scores. These were then normalized to the median function score of synonymous variants (exon 10 screen) or to the median function score of intronic variants (non-coding screen) in each condition, prior to averaging function scores across conditions. Function scores averaged across conditions were once more normalized to the median function score of synonymous variants (saturation screen) or intronic variants (non-coding screen), and final function scores were assigned from variants scored in at least two conditions.

A high-stringency pegRNA activity filter was used to re-analyze the *MLH1* exon 10 screen. This was set to the mean ST editing threshold above which the range of pLoF (i.e., nonsense and canonical splice) variant scores is non-overlapping with the range of pNeut (i.e., synonymous) variant scores, corresponding to 36% mean ST editing per variant.

To classify variants in each screen as LoF, a normal distribution of neutral function scores was modeled from synonymous variants (*SMARCB1* and *MLH1* exon 10 screens) or intronic variants (*MLH1* non-coding screen) to calculate p-values for each function score. To correct for multiple hypothesis testing, the Benjamini-Hochberg (BH) procedure was applied. Variants with q-values less than 0.05 for the *SMARCB1* screen and less than 0.01 for the *MLH1* screen were deemed LoF variants.

### AUC analysis

AUC measurements for separating pLoF and pNeut variants by function score were computed over a continuous range of ST editing thresholds. At each ST editing threshold sampled, function scores were re-calculated from the set of pegRNAs with ST editing percentages above threshold. For the *MLH1* exon 10 screen, synonymous variants were defined as pNeut and nonsense and canonical splice variants as pLoF. For the *MLH1* non-coding screen, intronic variants were defined as pNeut and canonical splice variants as pLoF.

To define a high-stringency ST editing threshold for the exon 10 screen, AUC measurements for distinguishing LoF variants by function score were computed over a continuous range of mean ST editing thresholds. Mean ST editing rates were calculated by averaging the editing rates of pegRNAs programming the same variant across conditions. The high-stringency analysis excluded pegRNAs with ST editing rates less than 5% in PEmax cell lines or less than 25% in PEmax-MLH1dn cell lines, as these were removed prior.

### Comparing function scores with orthologous data sets

The crystal structure of human MLH1 (PDB: 4P7A) was imported to PyMol. Maximum function scores for missense variants tested at each amino acid position were calculated and used to color-code residues. Negative function scores were set to 0.0 for this analysis.

*MLH1* variant pathogenicity assertions were retrieved from ClinVar in August 2023. For benchmarking of function scores, variants deemed “pathogenic” or “likely pathogenic” were grouped together, as were variants deemed “benign” or “likely benign”. Variants deemed to be of “uncertain significance” or with “conflicting interpretations of pathogenicity” were also grouped together as VUS.

CADD scores (v1.6)^50^ and annotations were obtained for all *SMARCB1* and *MLH1* SNVs (https://cadd.gs.washington.edu/download), inclusive of SpliceAI scores. A single maximum SpliceAI score was determined for each variant from independent scores for “acceptor gain”, “acceptor loss”, “donor gain”, and “donor loss” scores and used for comparison to function scores. Variant “consequence” annotations for MNVs, deletions, and insertions were manually curated. The variant c.-7_1del was annotated as a 5’-UTR variant throughout analyses except where explicitly indicated “start loss”.

## DATA AND CODE AVAILABILITY

All experimental data including pegRNA frequencies, pegRNA scores, ST editing rates, and function scores are available in Supplementary Information. Raw NGS data (.fastq files) will be deposited to a public archive and made available prior to publication.

Custom scripts used in this study are available on GitHub: https://github.com/FrancisCrickInstitute/PooledPEScreen.

## ACKNOWLEDGEMENTS

We thank the Crick’s Advanced Sequencing Scientific Technology Platform (STP) for performing sequencing and the Crick’s Cell Services STP for maintenance of cell lines.

This work was supported by the Francis Crick Institute which receives its core funding from Cancer Research UK (CC2190), the UK Medical Research Council (CC2190), and the Wellcome Trust (CC2190). For the purpose of Open Access, the authors have applied a CC BY public copyright license to any Author Accepted Manuscript version arising from this submission.

## AUTHOR CONTRIBUTIONS

M.H., C.M.K., and G.M.F. conceived the project and designed experiments; C.M.K. and M.H. performed experiments and analyzed the data; M.H., C.M.K., M.B., A.C., M.S., and G.M.F. performed initial optimizations and derived key reagents; G.M.F. supervised the project; and M.H., C.M.K., and G.M.F. wrote the manuscript with input from all authors.

## SUPPLEMENTAL INFORMATION

**Supplementary Table 1. Rates of ST editing by pegRNA scaffold library.**

**Supplementary Table 2. Rates of ST editing for comparison of SNVs and MNVs.**

**Supplementary Table 3. *SMARCB1* ST editing rates and pegRNA scores.**

**Supplementary Table 4. Function scores derived for variants in *SMARCB1*.**

**Supplementary Table 5. ST editing rates and pegRNA scores for the *MLH1* exon 10 screen.**

**Supplementary Table 6. Function scores for the *MLH1* exon 10 screen.**

**Supplementary Table 7. ST editing rates and pegRNA scores for the *MLH1* non-coding screen.**

**Supplementary Table 8. Function scores for the *MLH1* non-coding screen.**

**Supplementary Table 9. Oligonucleotides used in this study.**

